# The Cu(II) reductase RclA protects *Escherichia coli* against the combination of hypochlorous acid and intracellular copper

**DOI:** 10.1101/690669

**Authors:** Rhea M. Derke, Alexander J. Barron, Caitlin E. Billiot, Ivis F. Chaple, Suzanne E. Lapi, Nichole A. Broderick, Michael J. Gray

**Author notes:** Corresponding Author: Michael J. Gray, 656 Bevill Biomedical Research Building, 845 19th Street South, Birmingham, AL 35294.

## Abstract

Inflammatory bowel diseases (IBDs) are a growing health concern. Enterobacteria, including *Escherichia coli*, bloom to high levels in the gut during inflammation and strongly contribute to the pathology of IBDs. To survive in the inflamed gut, *E. coli* must tolerate high levels of antimicrobial compounds produced by the immune system, including toxic metals like copper and reactive chlorine oxidants like hypochlorous acid (HOCl). In this work, we show that the widely-conserved bacterial HOCl resistance enzyme RclA catalyzes the reduction of copper (II) to copper (I), and specifically protects *E. coli* against the combination of HOCl and intracellular copper, probably by preventing Cu(III) accumulation. *E. coli* lacking RclA were highly sensitive to HOCl and were defective in colonizing an animal host. Our results indicate unexpected complexity in the interactions between antimicrobial toxins produced by innate immune cells and suggest an important and previously unsuspected role for copper redox reactions during inflammation.

## INTRODUCTION

Inflammatory bowel diseases (IBDs), like Crohn’s disease and ulcerative colitis, are a growing health problem (1), and are associated with dramatic changes in the composition of the gut microbiome (2–6). Patients with IBDs have increased proportions of proteobacteria, especially *Escherichia coli* and other *Enterobacteriaceae*, in their gut microbiomes, which is thought to contribute to the progression of disease (7–10). The bloom of enterobacteria in the inflamed gut is driven by increased availability of respiratory terminal electron acceptors (*e.g.* oxygen, nitrate, TMAO) and carbon sources (*e.g.* ethanolamine, mucin), which *E. coli* and other facultative anaerobes can use to outcompete the obligate anaerobes (*Bacteroides* and *Clostridia*) that dominate a healthy gut microbiome (2, 3). In addition to these nutritional changes in the gut environment, inflammation also leads to infiltration of innate immune cells (*e.g.* neutrophils) into the lumen of the gut (11) and an associated increased production of antimicrobial compounds by the innate immune system (9, 10, 12), which also impact the bacterial community in the gut. These include antimicrobial peptides, toxic metals (*e.g.* copper), reactive oxygen species (ROS), reactive nitrogen species (RNS), and reactive chlorine species (RCS) (3, 13, 14). Since bacteria living in an inflamed gut are likely to be exposed to substantially increased levels of these toxins, the differential survival of proteobacteria during long-term inflammation suggests that *E. coli* may have evolved better mechanisms to resist these stresses than other types of commensal bacteria.

Innate immune cells use copper as an antimicrobial agent and copper levels rise in inflamed tissues, although the exact mechanism(s) by which copper kills bacteria are not yet fully understood (15–17). RCS are highly reactive oxidants produced by neutrophils and are potent antibacterial compounds (18, 19) that have been reported to play a role in controlling intestinal bacterial populations (20–22). *E. coli* does not efficiently survive being phagocytosed by neutrophils (23, 24), but the release of RCS-generating myeloperoxidase from neutrophils in the inflamed gut (11) means that *E. coli* is exposed to increased levels of RCS in that environment. The RCS response of *E. coli* is complex and incompletely understood (19, 25–28), but characteristically involves repair of damaged cellular components, often proteins (19). Protein-stabilizing chaperones are upregulated during RCS stress, including Hsp33 (29, 30) and inorganic polyphosphate (28, 31, 32), and enzymes are expressed that repair oxidized proteins, including periplasmic methionine sulfoxide reductase (MsrPQ) (33) and the chaperedoxin CnoX (34). *E.* coli has multiple HOCl-sensing regulators, including YedVW (which regulates MsrPQ) (33), NemR (regulator of NemA and GloA, which detoxify reactive aldehydes) (25), HypT (which regulates cysteine, methionine, and iron metabolism) (27), and RclR, the RCS-specific activator of the *rclABC* operon (26).

The *rclA* gene of *E. coli* encodes a predicted flavin-dependent oxidoreductase, is upregulated more than 100-fold in the presence of RCS (26, 35), and protects against killing by HOCl via a previously unknown mechanism (26). RclA is the most phylogenetically conserved protein of the Rcl system (Figure 1) and is found almost exclusively in bacteria known to colonize epithelial surfaces (Supplemental Table 1), suggesting that it may play an important role in host-microbe interactions in many species. Bacteria encoding RclA homologs include Gram negative species (*e.g. Salmonella enterica*), Gram positive species (*e.g. Streptococcus sanguinis*), obligate anaerobes (*e.g. Clostridium perfringens*), facultative anaerobes (*e.g. Staphylococcus aureus*), pathogens (*e.g. Enterococcus faecalis*), commensals (*e.g. Bacteroides thetaiotaomicron*), and probiotics (*e.g. Lactobacillus reuteri*) (36), suggesting that RclA’s function may be broadly conserved and not specific to a single niche or type of host-microbe interaction. RclR, RclB, and RclC are much less widely conserved, and are found only in certain species of proteobacteria, primarily members of the *Enterobacteriaceae* (Figure 1 and Supplemental Table 1).

**Figure 1:**
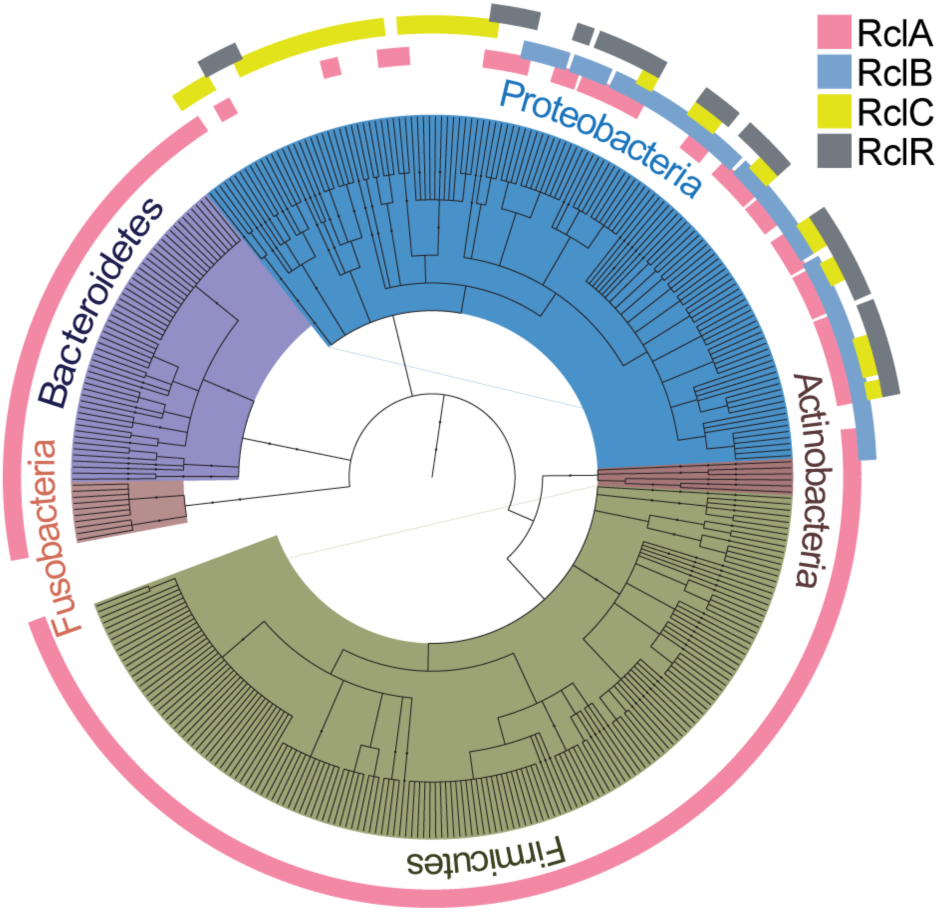
RclA is widely conserved among bacteria that colonize epithelial surfaces. Phylogenetic tree made from amino acid sequence alignments of RclA (284 species), RclB (61 species), RclC, (49 species), and RclR (43 species), relative to the respective proteins in *E. coli* MG1655. See Supplemental Data Table 1 for lists of each hit used in the phylogenetic tree. Tree graphic was made using the interactive tree of life (37).

In this work, we have now determined that RclA is a thermostable, HOCl-resistant copper (II) reductase that is required for efficient colonization of an animal host and protects *E. coli* specifically against the combination of HOCl and intracellular copper, probably by preventing the formation of highly reactive Cu(III). We also found that, surprisingly, extracellular copper effectively protects bacteria against killing by HOCl both in cell culture and in an animal colonization model, most likely by catalyzing the breakdown of HOCl before it reaches the bacterial cells. These findings reveal a previously unappreciated interaction between two key inflammatory antimicrobials and a novel way in which a commensal bacterium responds to and resists the combinatorial stress caused by copper and HOCl.

## RESULTS

### RclA contributes to HOCl resistance and host colonization

An rclA mutant of E. coli is more susceptible to HOCl-mediated killing than the wild-type (26). In this work, we utilized a growth curve-based method to measure sensitivity to sub-lethal HOCl stress by quantifying changes in lag phase extension (LPE) of cultures grown in the presence of HOCl. Using this method, which we have found to be considerably more reproducible than other techniques for assessing bacterial HOCl sensitivity, we observed significant increases in LPE for a ΔrclA mutant strain compared to the wild type grown in the presence of various concentrations of HOCl (Supplemental Figure 1). We observed similar trends when the assay was done on stationary- and log-phase cells (Supplemental Figure 2A and B), so stationary phase cultures were used for subsequent assays. Furthermore, we determined that there was no drop in CFU after treatment with HOCl at these concentrations, indicating that LPE measures recovery of cultures from non-lethal stress (Supplemental Figure 2C). These results confirm previous results and further illustrate the importance of RclA in resisting HOCl-mediated oxidative stress in E. coli.

To directly test the role of RclA in interactions with an animal host, we examined the ability of *E. coli* to colonize the intestine of *Drosophila melanogaster*, where the presence of enterobacteria is known to stimulate antimicrobial HOCl production by the dual oxidase Duox (20, 22). Since an *E. coli* K-12 strain did not efficiently colonize *D. melanogaster* (Supplemental Figure 3), we used the colonization-proficient *E. coli* strain Nissle 1917 (EcN) (38, 39) in these experiments. EcN Δ*rclA* mutants had a significant defect in their ability to colonize NP1-GAL4 *D. melanogaster* flies compared to wild-type EcN at 3- and 8-hours post infection (hpi) (Figure 2, filled circles). This shows that *rclA* is important for EcN to survive host responses during early colonization.

**Figure 2:**
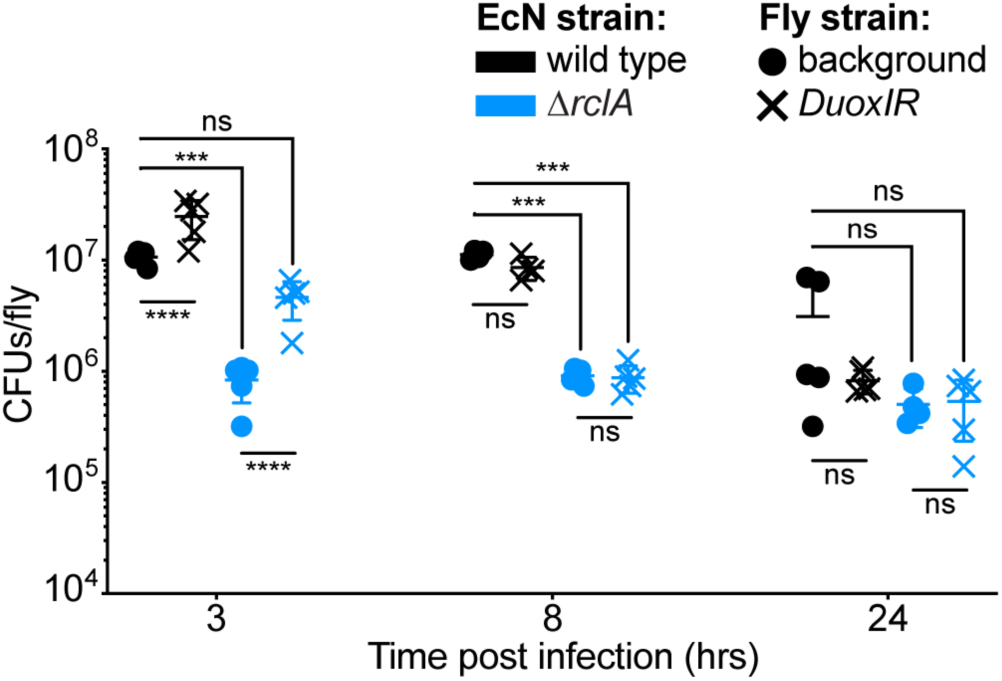
EcN lacking *rclA* colonize *D. melanogaster* less effectively and early colonization with *ΔrclA* EcN is improved in the absence of Duox-mediated oxidation. Flies were fed either wild-type or *ΔrclA* EcN (1×10^11^ CFU/mL) and bacterial loads were measured at the indicated times post infection (n = 4 - 5, ± SD). RCS-deficient Duox-RNAi flies were obtained from crosses of *UAS-dDuox-RNAi* with *NP1-GAL4* (enterocyte - specific driver). Statistical analysis was performed using a two-way ANOVA with Tukey’s multiple comparisons test (**** = P < 0.0001, *** = P < 0.001, ns = not significant).

To investigate the role of host-produced RCS in the colonization defect of the *rclA* EcN mutant, we reduced the gut-specific expression of Duox in the flies using Duox-RNAi and repeated the colonization experiments. Both strains colonized significantly better at 3 hpi when Duox was knocked down in the flies (Figure 2, x symbols), which was expected because *rclA* is not the only gene known to contribute to HOCl resistance in *E. coli* (19). Importantly, the colonization defect of *ΔrclA* EcN at 3 hpi was abrogated in the *DuoxIR* flies, with CFUs/fly not being significantly different from wild-type EcN colonizing flies that are able to express Duox in the gut (Figure 2). We confirmed that HOCl production was reduced in the *DuoxIR* flies using the HOCl-sensing fluorescent probe R19-S (Supplemental Figure 4) (40). Taken together, these results show that *rclA* facilitates colonization of an animal host during early timepoints and indicate that *rclA* relieves stress caused by host-produced oxidation.

### RclA is homologous to mercuric reductase (MerA) and copper response genes are upregulated after HOCl stress in EcN

Although the fact that *rclA* protects *E. coli* from RCS was previously known (26), the mechanism by which it does so was not. Based on its homology to other flavin-dependent disulfide oxidoreductases (41), we hypothesized that RclA catalyzed the reduction of an unknown cellular component oxidized by RCS. RclA is homologous to mercuric reductase (MerA), an enzyme that reduces Hg(II) to Hg(0) (42). These sequences are particularly well conserved at the known active site of MerA, a CXXXXC motif (Figure 3). However, MerA has an extra N-terminal domain and two additional conserved cysteine pairs used in metal binding (42) while RclA only has one conserved cysteine pair (Supplemental Figures 5 and 6). This indicates that if RclA does interact with metal(s), those interactions must be mediated through mechanisms different from those of MerA.

**Figure 3:**
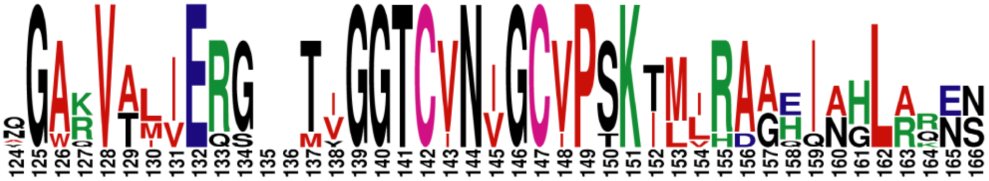
RclA and Mercuric reductase (MerA) share a conserved active site. Active site alignment of *E. coli* RclA and MerA amino acid sequences from seven bacterial species (*Escherichia coli*, *Staphylococcus aureus*, *Salmonella enterica*, *Listeria monocytogenes*, *Klebsiella pneumoniae*, *Serratia marcescens)*. Alignment was made using CLUSTAL O (1.2.4) and graphic was made with WEBLOGO.

HOCl-stressed *E. coli* K-12 downregulates genes encoding iron import systems (*e.g. fepABCD, fhuACDF*) and upregulates genes for zinc and copper resistance (*e.g. copA, cueO, cusC, zntA, zupT*) (25), also suggesting metals may play some role in RCS resistance. Transcriptomic profiling of EcN after treatment with a sublethal dose of HOCl confirmed this and revealed the upregulation of several genes encoding proteins involved in response to copper toxicity (Figure 4, Supplemental Table 2), despite the very low amounts of copper present in the media used in that experiment (9 nM) (43). These include members of the Cus and Cue export systems, which are factors appreciated for their role in preventing copper toxicity and importance for *E. coli* colonization within mammalian hosts (44–47). This suggested that copper might play an important role during HOCl stress for EcN. The homology between RclA and MerA and the indication that copper and HOCl responses may be connected in *E. coli* led us to investigate the role of *rclA* in resisting HOCl stress under growth conditions containing different amounts of copper.

**Figure 4:**
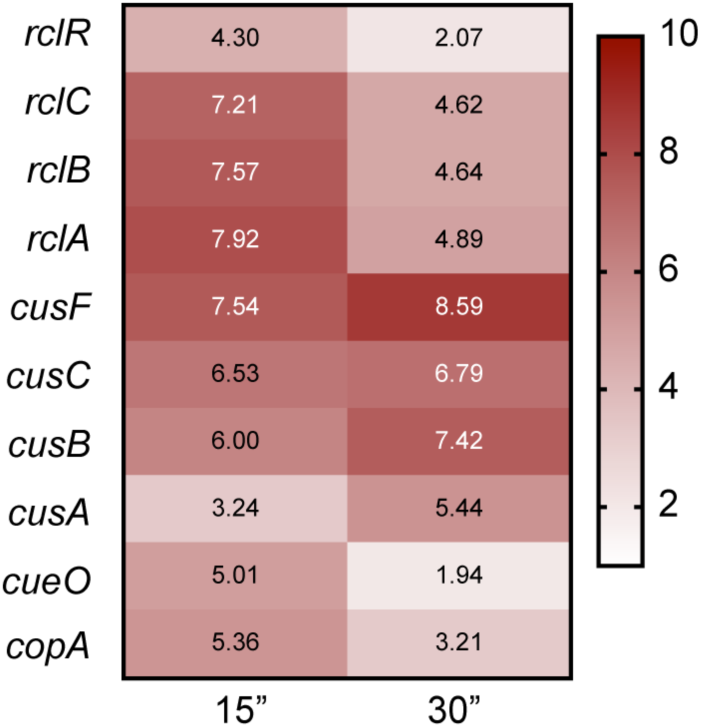
EcN upregulates copper export proteins during HOCl stress. Log_2_ fold-change in EcN gene expression at 15 and 30 minutes after non-lethal HOCl treatment (0.4 mM), relative to a control sample taken directly before treatment. mRNA counts were measured using RNA sequencing and analyzed with Bowtie2.

### Extracellular CuCl_2_ protects both wild-type and Δ*rclA E. coli* against HOCl

How the presence of copper influences bacterial sensitivity to RCS has not been investigated before this study. We first used growth curves in the presence of copper and HOCl to identify how combinations of HOCl and extracellular copper influenced the sensitivity of wild type and Δ*rclA* mutant *E. coli* (Figure 5A and 5B). As we observed above, the Δ*rclA* mutant was more sensitive to HOCl in MOPS, but the sensitivity to HOCl of both strains was greatly increased when Cu was removed from the media, indicating that the low concentration of Cu present in MOPS medium (9 nM) (43) was enough to react with the added HOCl and change the sensitivity of our strains. Consistent with this and the ability of Cu to catalyze the decomposition of HOCl to non-toxic O_2_ and Cl^-^ (48–51), addition of 10 µM CuCl_2_ to HOCl-containing media greatly decreased sensitivity of both the wild type and the *rclA* null mutant to HOCl (Figure 5). Copper being uniformly protective for both strains makes it likely that extracellular copper had reacted with and detoxified the HOCl before cells were inoculated into the media. To confirm this result, we tested if the addition of exogenous copper (0.5 mM CuCl_2_) could protect *E. coli* against killing by a very high concentration of HOCl (1 mM). We found that treatment with 1 mM HOCl resulted in complete killing of both strains (Figure 5C). Both strains survived several orders of magnitude better when 0.5 mM CuCl_2_ was added to the media immediately before HOCl stress (Figure 5C). In addition, colonization of flies by EcN was enhanced when their diet was supplemented with copper (Supplemental Figure 8), consistent with Cu detoxifying HOCl *in vivo*. Taken together, these results illustrate that the presence of exogenous copper strongly protects *E. coli* against HOCl.

**Figure 5:**
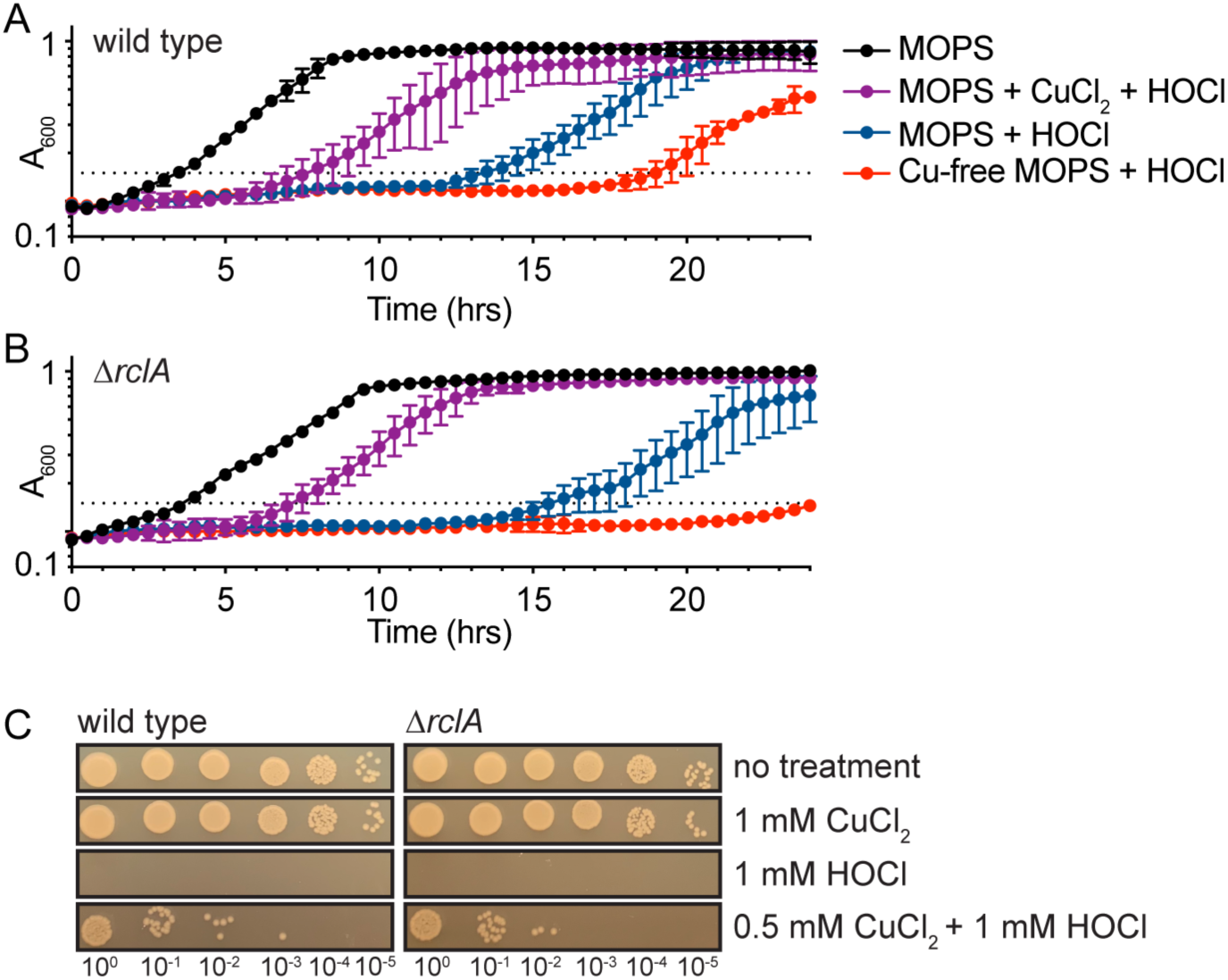
Extracellular CuCl_2_ protects both wild-type and Δ*rclA E. coli* against HOCl. Growth curves of wild type (A) and Δ*rclA* (B) strains in MOPS with the addition of various combinations of HOCl (132 µM) and CuCl_2_ (10 µM) (n = 3, ± SD for each treatment). Only one of several HOCl concentrations tested is shown for simplicity, see Supplemental Figure 7 for lag phase extension calculations and a summary of all the conditions tested using the growth curve method. (C) Exogenous copper protects both wild type and Δ*rclA E. coli* from lethal HOCl stress (10 min at 1 mM).

### RclA protects *E. coli* against the combination of HOCl and intracellular copper

Next, we aimed to investigate how intracellular copper affects the HOCl resistance of *E coli*. To address this, we grew wild type and Δ*rclA* mutant *E. coli* overnight in minimal media with or without copper before inoculating the strains into copper-free media to perform HOCl-stress growth curves. Growing overnight cultures in media lacking copper was expected to starve the cells for this metal, thereby reducing the concentration of intracellular copper in those cultures. A broad range of HOCl concentrations were assayed to account for quenching of the oxidant by media components.

Consistent with the results in Figure 5, *E. coli* was more sensitive to inhibition by HOCl in media without copper. The Δ*rclA* mutant was more sensitive to HOCl than the wild type when grown overnight in copper-containing media, but this phenotype was lost when cells were starved for copper before stress (Figure 6), indicating that the physiological role of RclA is to resist the stress resulting from the combination of HOCl and copper in the cytoplasm.

**Figure 6:**
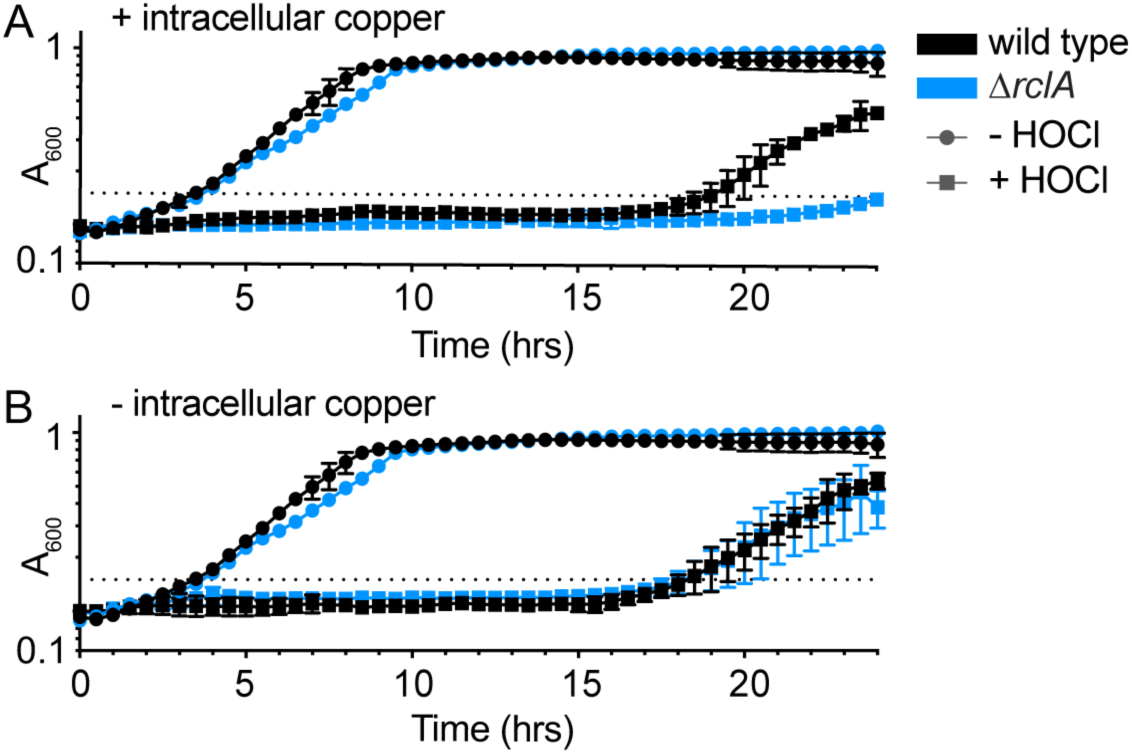
*rclA* is required to resist the combination of HOCl and intracellular Cu. HOCl (132 µM) growth curves (n = 3, ± SD) in copper-free MOPS of wild-type and Δ*rclA E. coli* after being grown up overnight in MOPS with (A) and without copper (B). Copper-free MOPS was prepared by treating the media with Chelex-100™ Chelating Resin (Bio-Rad 1421253) and adding back all the metals (metal stock solutions prepared in metal-free water, Optima™ LC/MS Grade, Fisher Chemical W6-1), except for copper, to the published concentrations (43). In cells containing intracellular copper (A), the Δ*rclA* mutant has delayed growth relative to the wild type. There is no difference between the strains when the cells are starved for intracellular copper (B). One HOCl concentration is shown for simplicity, see Supplemental Figure 9 for growth curves showing more HOCl concentrations.

### RclA reduces copper (II) to copper (I)

Based on the effect of copper starvation on the HOCl sensitivity of the Δ*rclA* mutant, the sequence homology between RclA and MerA, and the predicted oxidoreductase activity of RclA (41), we hypothesized that the substrate of RclA might be copper. The reaction between copper and HOCl is known to generate strong oxidizing intermediates, most likely highly reactive Cu(III) (48–52). HOCl is also capable of oxidizing other transition metals, including iron (53, 54) and manganese (55, 56). We therefore measured the specific activity (SA) of purified RclA in the presence of a panel of biologically relevant metals. We also included mercury in the panel of metals because of RclA’s homology to MerA, although it is unlikely to be physiologically relevant since we do not expect *E. coli* to encounter this metal in its environment under normal conditions. We note that the oxidized forms of many transition metals are insoluble in aqueous solution, which limited the set of substrates we could test with this experiment.

In the absence of any metal, RclA slowly oxidized NADH (0.0303 µmole NAD^+^ min^-1^ mg^-1^ RclA), consistent with the background activity of other flavin-dependent oxidoreductases in the absence of their specific substrates (57, 58). Three of the metals we tested significantly affected RclA SA, as measured by NADH oxidation. Copper and mercury both significantly increased the SA of RclA, while zinc caused a decrease in SA (Figure 7A). Copper is a potent inhibitor of MerA activity (59), further emphasizing the distinct nature of these two enzymes. RclA oxidized NADPH at similar rates to NADH in the absence of metals, but there was no significant increase to SA when copper was added to the reactions. The addition of exogenous thiols, commonly added as β-mercaptoethanol (BME), is required for MerA activity (60). To determine if exogenous thiols increase the reaction rate of RclA, we added 1 mM BME to the RclA reactions. BME rapidly reduced Cu(II) to Cu(I) in the absence of RclA, but had no effect on the SA of NADH reduction by RclA with or without the addition of copper (Supplemental Figure 10).

**Figure 7:**
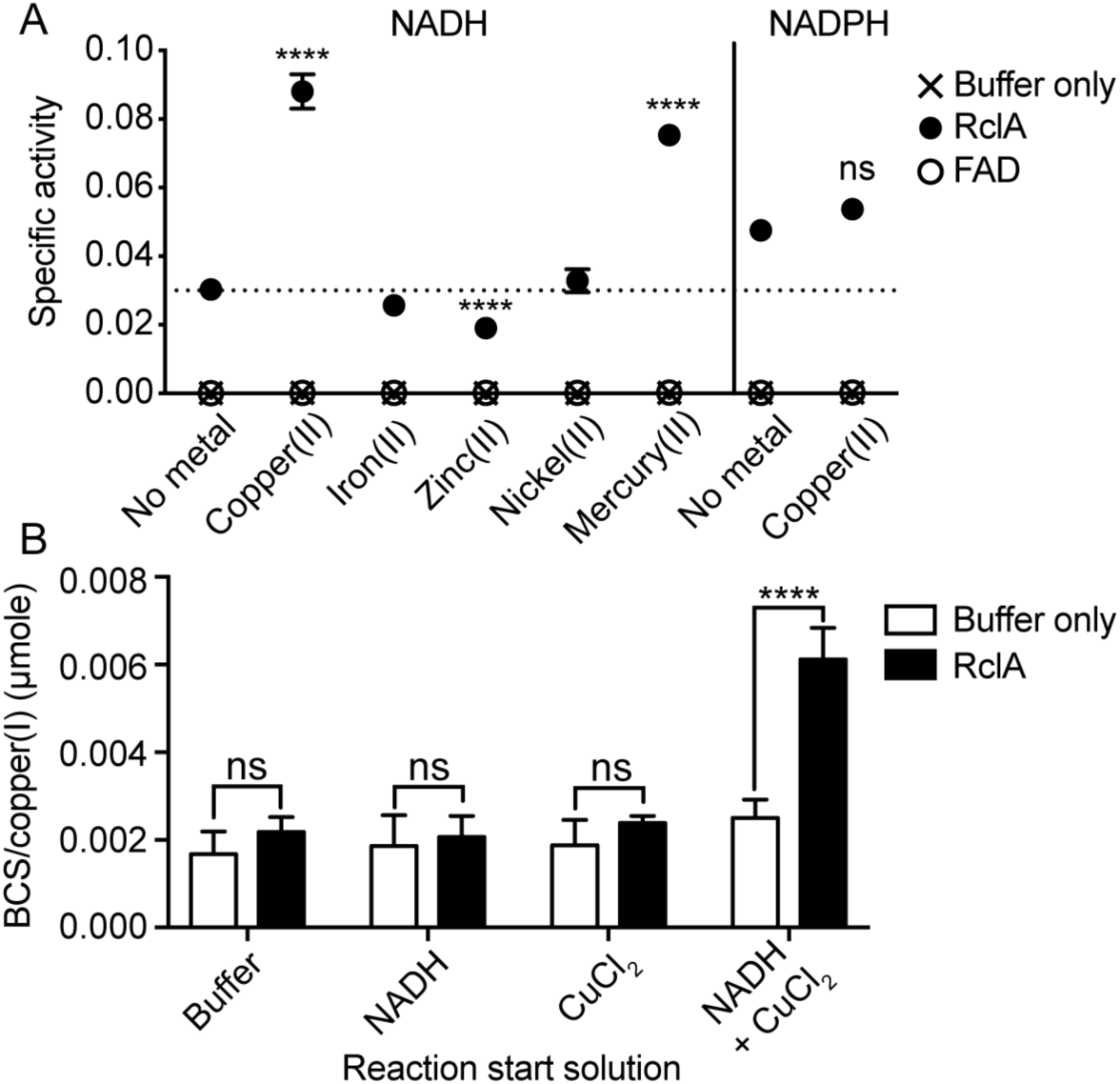
RclA reduces Cu(II) to Cu(I). (A) RclA specific activity (SA) increases in the presence of Cu(II) and Hg(II). SA (µmole NAD(P)^+^ min^-1^ mg^-1^ RclA) of RclA was assayed by measuring NAD(P)H oxidation over time spectrophotometrically (n = 6, ± SD).

Since RclA is an NADH oxidase, the results shown in Figure 7A strongly suggested that this enzyme was concurrently reducing copper. Copper exists in four possible oxidation states, Cu(I), Cu(II), and the less common and highly reactive Cu(III) and Cu(IV) states (61). The copper salt used in our RclA SA determinations was CuCl_2_, suggesting that RclA was reducing this Cu(II) species to Cu(I). This was very surprising, since Cu(I) is generally thought of as a toxic species that causes oxidative stress (46, 62), so it was not obvious either why *E. coli* would produce Cu(I) during RCS stress or why production of Cu(I) might be protective. We therefore first sought to validate that RclA was in fact reducing Cu(II) to Cu(I) while oxidizing NADH to NAD^+^. We measured Cu(I) accumulation in RclA reactions directly using the Cu(I)-specific chelator bathocuproinedisulfonic acid (BCS) (63). NADH spontaneously reduces Cu(II) (64) at rates too slow to impact the measurements made here, but BCS increases the rate of this non-enzymatic copper reduction by shifting the equilibrium of the reaction towards Cu(I) (65). Stopping RclA reactions with a mixture of BCS and EDTA, to chelate any remaining Cu(II), allowed us to observe RclA-dependent Cu(I) accumulation (Supplemental Figure 11). We observed a significant increase in BCS/Cu(I) complex formation only in reactions containing RclA, NADH, and Cu(II), and not in reactions lacking any single component (Figure 7B, Supplemental Figure 11). Furthermore, we validated that the copper reductase activity of RclA was maintained when the reactions were performed anaerobically (Supplemental Figure 12) and that FAD alone did not catalyze Cu(II) reduction (Figure 7A). Taken together, these findings show that RclA has Cu(II) reductase activity and generates Cu(I) as a product.

Differences in SA in the presence of each metal were analyzed using a two-way ANOVA with Dunnet’s multiple comparison test using the no metal reaction as the control (****= P < 0.0001, ** = P < 0.01). (B) Cu(I) accumulates after the RclA and NADH/copper (II) reaction, as measured by BCS/Cu(I)-complex absorption. RclA reactions were started with the indicated solutions and carried out as described in A (n = 6, ± SD). Differences in the amount of BCS/Cu(I) complex between the buffer only and RclA reactions were analyzed using a two-way ANOVA with Dunnet’s multiple comparison test using the buffer only sample as the control for each reaction start solution (****= P < 0.0001, ns = not significant).

### RclA does not protect against HOCl by facilitating copper export

Since the copper exporters of *E. coli* (*copA* and *cusCFBA*) are upregulated by HOCl treatment (Figure 3, Supplemental Table 2) (25) and only transport Cu(I) (46, 47, 66, 67), it was possible that RclA facilitates the rapid export of cytoplasmic copper, allowing it to react with and eliminate HOCl outside the cell. To test whether the Cu(II) reductase activity of RclA is important for exporting copper during HOCl stress, we measured intracellular copper concentrations before and after HOCl stress with ICP mass spectrometry. We found that the Δ*rclA* mutant contained, on average, more intracellular copper before HOCl stress than the wild-type but that both strains contained similar amounts of copper after HOCl stress (Supplemental Figure 13). We constructed mutants lacking *copA*, which is reported to result in increased intracellular copper (47), but unexpectedly found that the *copA rclA* double mutant had a substantial growth defect in copper-free media, even in the absence of HOCl (Supplemental Figure 14). We do not yet know the explanation for this intriguing result. Overall, our results indicate that although RclA may affect basal copper homeostasis, it does not appear to protect *E. coli* against HOCl by facilitating export of copper at levels detectable using these methods.

### RclA is thermostable and resistant to denaturation by HOCl and urea

We hypothesized that the copper reductase activity of RclA was likely to be relatively stable under denaturing conditions because it must remain active during exposure to HOCl stress, which is known to cause extensive protein misfolding and aggregation *in vivo* (19, 27, 28, 32–34). To test this hypothesis, we first measured RclA activity after treatment with protein denaturing agents (HOCl and urea) *in vitro*. HOCl treatment (with 0, 5, 10, and 20-fold molar ratios of HOCl to RclA) was done on ice for 30 minutes and urea treatment (0, 2, 4, and 6 M) was carried out at room temperature for 24 hours. RclA retained full copper reductase activity at all HOCl levels tested, indicating that it is highly resistant to treatment with HOCl (Figure 8A). By comparison, the NADH oxidase activity of lactate dehydrogenase was significantly decreased after treatment with a 5-fold excess of HOCl (Figure 8B). RclA also retained a remarkable 35.8% of full activity after being equilibrated in 6 M urea (Figure 8C). Finally, we used circular dichroism (CD) spectroscopy to measure the melting temperature (T_m_) of RclA, which was 65 °C (Figure 8D, Supplemental Figure 15), indicating that RclA is thermostable relative to the rest of the *E. coli* proteome, which has an average T_m_ of 55 °C (standard deviation = 5.4°C) (49, 68).

**Figure 8.**
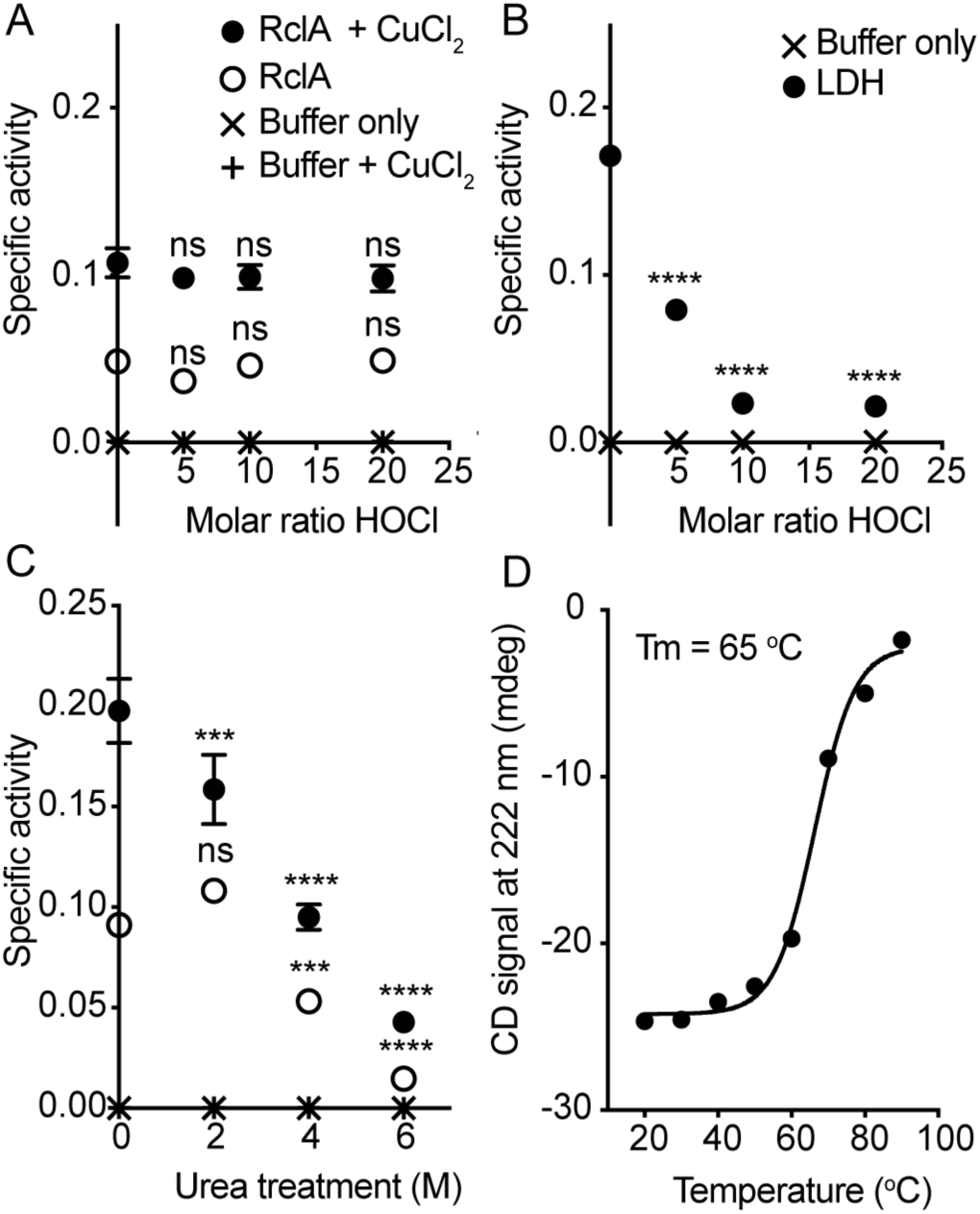
RclA is resistant to denaturation. (A) SA of RclA (µmoles NAD^+^ min^-1^ mg RclA^-1^), with and without CuCl_2_, after being treated with the indicated molar ratios of HOCl to RclA. (B) SA of lactate dehydrogenase (LDH) (µmoles NAD^+^ min^-1^ U LDH^-1^) used as control reactions for HOCl degradation of enzymatic activity. (C) RclA with and without CuCl_2_ (200 µM final) after being treated with the indicated concentrations of urea. (D) CD signals at 222 nm (mdeg) (raw data shown in Supplemental Figure 15) at each temperature used to determine the T_m_ of RclA (65 ^°^C). Differences in SA (n = 6, ± SD) after treatment were analyzed using a two-way ANOVA with Sidak’s multiple comparison test for HOCl treatment (A and B) and Dunnet’s test using the buffer only reaction as the control for the urea treated samples (C) (**** = P < 0.0001, * = P < 0.05 ns = not significant).

## DISCUSSION

The antimicrobial function of copper in host-microbe interactions is well-established (15, 17, 44, 69), although the exact mechanism(s) by which copper kills bacteria remain incompletely known (70, 71). In this work, we identified a new way in which copper toxicity contributes to host-bacteria interactions via its reactions with RCS. We identified RclA as a highly stable Cu(II) reductase (Figure 7) required for resisting killing by the combination of HOCl and intracellular copper in *E. coli* (Figure 6). In the absence of *rclA*, *E. coli* had a significant defect in the early stages of host colonization which was partially eliminated when production of HOCl by the host was reduced (Figure 2). Duox activation and HOCl production are rapid host responses that occur at early stages of bacterial colonization of the gut (20), in agreement with our data showing that RclA is important for initial establishment. As the course of infection progresses, additional antimicrobial effectors, such as antimicrobial peptides regulated by NF-κB signaling, become more prominent (27), which could explain the similar colonization levels of mutant and wild-type strains at later time points. The amount of copper in bacterial cells is low (15, 16), but how much is unbound by protein and its redox state under different conditions are unknown (15). Given the broad conservation of RclA among host-associated microbes, we propose that there is likely to be a common and previously unsuspected role for copper redox reactions in interactions between bacteria and the innate immune system.

Copper accumulates in host tissues during inflammation (72, 73), as do RCS (74, 75). Our discovery that even very low concentrations of extracellular copper can protect bacteria against RCS both *in vitro* and *in vivo* adds a new and important facet to understanding copper’s role in innate immunity. Since a large proportion of host tissue damage during inflammation is due to HOCl (76, 77), the presence of copper in inflamed tissues may play an important role not only in killing bacteria, but potentially in protecting host cells, although this hypothesis will require further testing. Our results also show that media copper concentrations are a key variable in experiments testing the sensitivity of cells to HOCl, and that care must be taken to account for media copper content and use metal-free culture vessels in such experiments.

Both HOCl and copper cause oxidative stress in bacteria and Cu(I) is generally considered more toxic than Cu(II) (15, 19, 44, 69, 70, 78), so we were surprised that a Cu(II) reductase protected *E. coli* against HOCl. Copper reacts with the ROS hydrogen peroxide (H_2_O_2_) to form highly reactive hydroxyl radicals (17, 45, 62, 71), but how the presence of copper influences bacterial sensitivity to RCS has not been investigated before this study. The chemistry of reactions between HOCl and copper is complicated. HOCl can oxidize Cu(II) to highly reactive Cu(III) (48–52) and both Cu(I) and Cu(II) are known to catalyze the breakdown of HOCl (48–51). At near-neutral pH, similar to that in the large intestine or bacterial cytoplasm, Cu(I) accelerates the rate of decomposition of HOCl to O_2_ and chloride ions by as much as 10^8^-fold (49). One possibility to explain the protective effect of RclA is that it might facilitate an HOCl-degrading Cu(I) / Cu(II) redox cycle in the cytoplasm. If this were the case, however, the Δ*rclA* mutant would become more sensitive to HOCl in the absence of copper, the opposite of what we actually observed (Figure 6). We therefore propose that RclA-catalyzed reduction of Cu(II) to Cu(I) likely acts to limit the production of Cu(III) in the cytoplasm. Uncontrolled production of Cu(III) could greatly potentiate the ability of HOCl to kill bacterial cells.

While RclA itself is widely conserved, the *rclABCR* locus as a whole is restricted to certain enteric proteobacteria, including *E. coli*, *Salmonella*, *Citrobacter*, *Raoultella*, *Serratia*, and *Shigella*. These genera are notable for their close association with gut inflammation and the ability of pathogenic strains to bloom to very high levels in the gut in disease states (2, 3, 7–10, 79, 80). We hypothesize that the ability to survive increased levels of antimicrobial compounds (including RCS) in the inflamed gut is important for the ability of enterobacteria to exploit this niche, and our *in vivo* results with the Δ*rclA* mutant support this idea (Figure 2). The rate at which RclA oxidized NADH in the presence of copper *in vitro* was slow (approximately 4.4 min^-1^) (Figure 7), suggesting that we have not yet identified optimal reaction conditions for this enzyme. However, expression of *rclA* is rapidly induced >100-fold after sublethal doses of HOCl in *E. coli* (Figure 3, Supplemental Table 2) (26), which could compensate *in vivo* for the slow rate of NADH turnover we observed *in vitro*. We also do not currently know the physiological roles of RclB, which is a small predicted periplasmic protein, or RclC, which is a predicted inner membrane protein, although deletion of either of these genes results in increased HOCl sensitivity in *E. coli* (26). We hypothesize that they may form a complex with RclA *in vivo* and enhance its copper-dependent activity and are currently pursuing experiments to test this idea.

## EXPERIMENTAL PROCEDURES

### Strain and plasmid construction

*E. coli* strain MJG0586 (F^-^, λ^-^, *rph-1 ilvG^-^ rfb-50* Δ*rclA* λ(DE3 [*lacI lacUV5*-T7 gene 1 *ind1 sam7 nin5*])) was generated from MJG0046 (F^-^, λ^-^, *rph-1 ilvG^-^ rfb-50* Δ*rclA*) (26) using the Novagen DE3 lysogenization kit, according to the manufacturer’s instructions. The RclA coding sequence (1374 bp) was amplified from *E. coli* MG1655 genomic DNA with primers 5’ CTC GGT CTC CAA TGA ATA AAT ATC AGG CAG TGA 3’ and 5’ CTC GGT CTC AGC GCT TTA TTT GAC TAA TGA AAA TAG ATC A 3’ and cloned into the *Eco*31I (*Bsa*I) sites of plasmid pPR-IBA101 (IBA Lifesciences) to yield plasmid pRCLA10. pRCLA11 was generated by using QuikChange site directed mutagenesis (Agilent), modified to use a single mutagenic primer (5’ CTA TTT TCA TTA GTC AAA AGC GCT TGG AGC CAC CC 3’), to remove the stop codon between the RclA coding sequence and the twin-strep tag sequence in pRCLA10. *E. coli* Nissle 1917 Δ*rclA*::*cat^+^* strain MJG0846 was made by recombineering (81) using primers 5’ CGT CTA TAG TCA TGA TGT CAA ATG AAC GCG TTT CGA CAG GAA ATC ATC ATG GTG TAG GCT GGA GCT GCT TC 3’ and 5’ CTT TTC TCT GAG ACG CCA GAA TAT TTG TTC TGG CGT CTG ATT TTG AGT TTA CAT ATG AAT ATC CTC CTT AG 3’ to amplify the chloramphenicol resistance cassette from pKD3. The *cat*^+^ insertion was resolved using pCP20 resulting in the *E. coli* Nissle 1917 Δ*rclA* strain MJG0860.

### Protein expression and purification

Expression of twin-strep tagged RclA was done in MJG0586 containing pRCLA11 (MJG1338) in M9 minimal media (82) containing 2 g l^-1^ glucose and 100 µg ml^-1^ ampicillin. Overnight cultures of MJG1338 were diluted 1:100 and grown up to A_600_ = 0.4 at 37 ^°^C with shaking. When A_600_ = 0.4 was reached, expression was induced with IPTG (1 mM final concentration) and allowed to continue for 12-18 hours at 20 ^°^C. Purification of recombinant RclA was achieved to high purity using a 1 mL StrepTrap HP column (GE, 28-9136-30 AC) according to the manufacturer’s instructions. Purified protein was subsequently saturated with FAD cofactor by incubating the protein preparation with a 10-fold molar excess of FAD at room temperature for 45 mins. Excess FAD and elution buffer were dialyzed away with three exchanges of 1 L of RclA storage buffer (50 mM Tris-HCl, pH 7.5, 0.5 M NaCl, 2 mM DTT, 10% glycerol) at 4 ^°^C overnight.

### Colonization of D. melanogaster with E. coli Nissle 1917

*D. melanogaster stocks and husbandry: Canton-S* flies were used as a wild-type line. Duox-RNAi flies were obtained from crosses of *UAS-dDuox-RNAi* with *NP1-GAL4* (gut-specific driver). Unless otherwise noted, flies were reared on cornmeal medium (per liter of medium: 50g yeast, 70g cornmeal, 6g agar, 40g sucrose, 1.25ml methyl-paraben, and 5ml 95% ethanol). *Oral infections:* Adult female flies were starved for 2h at 29°C prior to being fed a 1:1 suspension of bacteria (OD_600_=200) and 2.5% sucrose applied to a filter paper disk on the surface of normal fly food. Suspensions of bacterial at OD_600_=200 were consistently found to contain 1×10^11^ CFU/mL (Supplemental Figure 16). A negative control was prepared by substituting LB for the bacterial suspension. Infections were maintained at 29°C. *CFU determination:* Flies were surface-sterilized in 70% ethanol and rinsed in sterile PBS. Individual flies were homogenized in screw-top bead tubes containing 600µl PBS. Dilution series of homogenates were prepared with a range of (10^0^ to 10^-5^). 3µl spots of each dilution were applied to LB plates and grown overnight at 30°C. *E. coli* colonies were identified by morphology and counted to determine CFUs/fly.

### RNA sequencing

*E. coli* Nissle 1917 was grown anaerobically (90% N_2_, 5% CO_2_, 5% H_2_, maintained with a Coy Laboratory Products anaerobic chamber) at 37°C in MOPS minimal media (Teknova) in sealed Hungate tubes to an A_600_ = 0.25, then treated anaerobically with 0.4 mM HOCl. Samples (7 ml) were harvested into 7 ml ice-cold isopropanol immediately before and 15 or 30 minutes after HOCl addition, harvested by centrifugation, and stored at −80°C until use. RNA was purified using the RiboPure™-Bacteria Kit (Ambion), according to the manufacturer’s instructions. mRNA-sequencing was performed at the UAB Heflin Center for Genomic Sciences, using an Illumina NextSeq 500 as described by the manufacturer (Illumina, Inc). The Agilent SureSelect Strand Specific mRNA library kit (Agilent) and ribosome reduction with the RiboMinus protocol for Gram-negative and Gram-positive bacteria (Life Technologies) were used as described by the manufacturer. The resulting mRNA was randomly fragmented with cations and heat, followed by first strand synthesis using random primers. Second strand cDNA production was done with standard techniques, and cDNA libraries were quantitated using qPCR in a Roche LightCycler 480 with the Kapa Biosystems kit for Illumina library quantitation (Kapa Biosystems) prior to cluster generation, which was performed according to the manufacturers recommendations for onboard clustering (Illumina). We generated approximately 8 million double-stranded 50 bp reads per sample. Data were analyzed using Bowtie2 and DEseq2. RNA sequencing data have been deposited in NCBI’s Gene Expression Omnibus (83) and are accessible through GEO Series accession number GSE144068.

### Growth curves for measuring sensitivity to HOCl

The molar HOCl concentration from concentrated sodium hypochlorite (Sigma Aldrich 425044) was quantified by measuring A_292_ of the stock solution diluted in 10 mM NaOH. Copper free MOPS was prepared by removing total metals from MOPS minimal media (Teknova) containing 2 g l^-1^ glucose and 1.32 mM K_2_HPO_4_ by treating prepared media with a universal chelator (Chelex® 100 Chelating Resin, Bio-Rad 1421253), filtering sterilizing away the Chelex resin, and adding back the metals normally present in MOPS suspended in metal-free water, except for copper. Overnight cultures of MG1655 and MJG0046 were grown up overnight in MOPS with and without copper. The overnight cultures were normalized to A_600_ = 0.8 in and diluted to A_600_ = 0.08 in MOPS minimal medium without copper with the indicated combinations/concentrations of HOCl and CuCl_2_. Cultures were incubated with shaking at 37 °C in a Tecan Infinite M1000 plate reader with A_600_ being measured every 30 minutes for 27 hours. Sensitivity was subsequently determined by comparing lag phase extensions (difference in hours to reach A_600_ ≥ 0.15 from the no HOCl treatment control under the same CuCl_2_ condition) for each stress condition.

### Determining the effect of copper on lethal HOCl stress

Wild type and Δ*rclA E. coli* MG1655 were grown overnight in MOPS minimal media. 500 µl of the overnight culture was pelleted, resuspended in copper-free MOPS, and subcultured into 9.5 ml of copper-free MOPS, then grown with shaking at 37°C to mid-log phase (A_600_ = 0.3 - 0.4). Once mid-log phase was reached, 1 mL of cells was aliquoted into microcentrifuge tubes for each treatment. Cells were treated by adding the indicated amounts of copper (II) chloride, followed by 1 mM HOCl, then incubated on a 37°C heat block for 10 minutes. After incubation, treatments were serially diluted in PBS and 5 µl aliquots were spotted on LB agar plates. Plates were dried and incubated at room temperature for 2 days.

### Measuring intracellular copper after HOCl stress

Wild-type and Δ*rclA E. coli* were grown up to A_600_ = 0.6 in MOPS and stressed with 400 µM HOCl for 30 mins at 37 °C with shaking. After 30 mins, the cultures were diluted 2-fold with MOPs to quench HOCl and pelleted. The mass of pellets was determined after being rinsed 3 times with PBS. Amount of copper was measured via ICP-MS (Agilent Technologies 7700x ICP-MS, Santa Clara, CA, USA) after suspending the collected pellets in concentrated nitric acid and diluting the suspensions to a 2% nitric acid matrix. Samples were filtered through 0.22 µM polytetrafluoroethylene filters to remove any particulates before running performing metal determination. Copper concentrations were determined by comparison to a standard curve (Agilent, 5188-6525) as calculated by Agilent software (ICP-MS MassHunter v4.3). Values determined by ICP-MS were normalized to pellet mass and dilution factor.

### NADH oxidase activity

Activity of purified RclA was assayed in 20 mM HEPES, 100 mM NaCl, pH 7 by measuring NADH oxidation over time spectrophotometrically. Purified recombinant RclA was preloaded into wells of a 96 well plate (100 µL of 6 uM RclA). Reactions were started by adding 100 µL of NADH (200 µM final) with the indicated metal salts (200 µM final) to RclA (3 µM final) at 37 °C. The absorbance of NADH (A_340_) was measured kinetically each minute for five minutes in a Tecan Infinite M1000 plate reader. A_340_ values were then used to calculate µmoles of NADH (ε_340_ = 6300 M^-1^ cm^-1^) and NADPH (ε_340_ = 6220 M^-1^ cm^-1^) at each timepoint. Absolute values of slopes of NAD(P)H (µmoles) over time (min) were divided by mg of RclA to determine the reported specific activities (SA, µmole NAD(P)^+^ min^-1^ mg^-1^ RclA) for each condition.

### Copper (I) quantification

Cu(I) accumulation after the course of the RclA reaction was measured using bathocuproinedisulfonic acid disodium salt (BCS, Sigma Aldrich B1125). NADH oxidation reactions were carried out as in the previous section but were started with either NADH (200 µM final), CuCl_2_ (200 µM final), NADH and CuCl_2_ (both at 200 µM final), or reaction buffer as a negative control to observe levels of background copper in the reagents used. Each reaction was stopped at 5 minutes with a BCS (400 µM final) and EDTA (1 mM final) solution using the injector system of a Tecan Infinite M1000 plate reader. The stopped reactions were incubated at 37 °C with the absorbance of BCS/Cu(I) complex being measured at 483 nm every minute for five minutes to ensure complete saturation of the BCS. The amount of the BCS/Cu(I) complex after 5 minutes was determined using the molar extinction coefficient 13,000 M^-1^ cm^-1^ (63).

### NADH oxidase activity after treatment with urea and HOCl

SA of RclA after being treated with denaturing agents was done by determining SA as above after incubation of RclA with either urea or HOCl. Urea treatment was done on 3 µM RclA for 24 hours at room temperature with increasing concentrations of urea (0, 2, 4, and 6 M) in reaction buffer (20 mM HEPES, 100 mM NaCl, pH 7). HOCl treatment was done by mixing increasing molar ratios of HOCl (0, 5, 10, and 20 X) with concentrated RclA or L - Lactate Dehydrogenase (LDH, Sigma LLDH-RO 10127230001) (35 µM) in oxidation buffer (50 mM sodium phosphate, pH 6.8, 150 mM NaCl) and incubating mixtures on ice for 30 minutes. HOCl treatment was quenched after 30 minutes by diluting the RclA solutions to 6 µM and the LDH solutions to 1 µM in reaction buffer (20 mM HEPES, 100 mM NaCl, pH 7). NADH oxidation reactions using LDH were performed under the same conditions as the RclA reactions but started with 1.2 mM NADH and 1.2 mM pyruvate.

### Melting temperature determination

Circular Dichroism (CD) spectra were obtained on a Jasco J815 circular dichroism spectrometer. CD spectra were collected on purified recombinant RclA exchanged into 20 mM HEPES, 100 mM NaCl, pH 7.5. Room temperature CD spectra in the range of 260 nm to 190 nm were obtained in 0.1 mm demountable quartz cells. Thermal CD data between 30 °C and 90 °C were obtained in standard 1.0 mm quartz cells. All data were collected with a 1.0 nm step size, 8 second averaging time per point, and a 2 nm bandwidth. Data were baseline corrected against the appropriate buffer solution and smoothed with Jasco software.

### Data analysis and bioinformatics

*Statistics*: All statistical analyses were performed using GraphPad Prism (version 7.0a). *Phylogenetic tree*: RclA conservation tree was made from amino acid sequence alignments of RclA (BLAST e value < 1×10^-90^ in 284 species), RclB (BLAST e value < 1×10^-1^ in 61 species), RclC, (BLAST e value < 1×10^-80^ in 49 species), and RclR (BLAST e value < 1×10^-40^ in 43 species). BLAST searches were done by comparing to each respective protein in *E. coli* MG1655. Tree graphic was made using the interactive tree of life (37) (https://itol.embl.de/). *Amino acid sequence alignments*: Active site alignment of *E. coli* RclA (ADC80840.1) and MerA amino acid sequences from seven bacterial species (*Escherichia coli*, ADC80840.1; *Staphylococcus aureus*, AKA87329.1; *Salmonella enterica*, ABQ57371.1; *Listeria monocytogenes*, PDA94520.1; *Klebsiella pneumoniae*, ABY75610.1; *Serratia marcescens*, ADM52740.1). Alignment was made using CLUSTAL O (1.2.4) (www.ebi.ac.uk/Tools/msa/clustalo/) and graphic was made with WEBLOGO (weblogo.berkeley.edu/logo.cgi). Full length alignment of *E. coli* RclA (ADC80840.1) and MerA (ADC80840.1). Alignment, conservation scoring, and graphic were made using PRALINE (www.ibi.vu.nl/programs/pralinewww/).

## ACKNOWLEDGEMENTS

Thanks to Dr. Peter Prevelige (UAB Department of Microbiology) for helpful discussions, Emily Schwessinger (University of Michigan) for preliminary characterization of RclA, Dr. Michael Jablonsky (UAB Chemistry Department) for assistance with CD spectra acquisition, Dr. Won-Jae Lee (Seoul National University) for providing the *D. melanogaster* Duox RNAi line, and William Van Der Pol and Ranjit Kumar at the UAB Center for Clinical and Translational Science for their help with the RNA sequencing analysis. This project was supported by University of Alabama at Birmingham Department of Microbiology startup funds, NIH grant R35GM124590 (to M.J.G), NIH grant R35GM128871 (to N.A.B.), and NIH training grant T90DE022736 (to R.M.D).

## AUTHOR CONTRIBUTIONS

R.M.D., A.J.B., C.E.B., and I.F.C. performed research, and R.M.D., A.J.B, S.E.L., N.A.B, and M.J.G. contributed to designing research, analyzing data, and writing the paper.

## SUPPLEMENTAL FIGURES

**Supplemental Figure 1:**
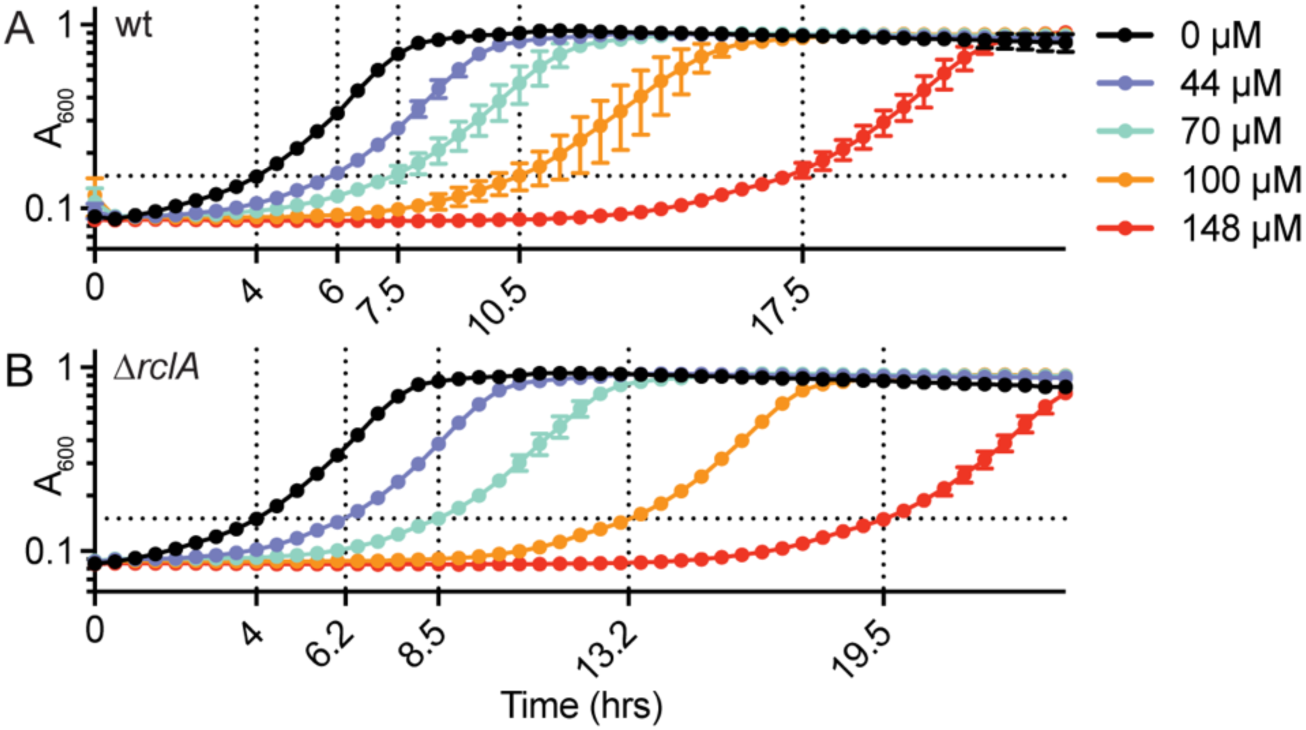
Growth curve method to determine HOCl sensitivity by measuring lag-phase extension. Overnight cultures of (A) MG1655 wild-type and (B) MJG0046 (Δ*rclA*) were normalized to A_600_ = 0.8 in MOPS minimal media and diluted to A_600_ = 0.08 in MOPS ± HOCl at the indicated concentrations. Cultures were incubated with shaking at 37 °C in a Tecan Infinite M1000 plate reader with A_600_ being measured every hour for 24 hours (n = 3, ± SD). Sensitivity was assessed by comparing lag phase extensions (difference in hours to reach A_600_ ≥ 0.15 from no HOCl control; vertical dotted lines) for each HOCl concentration.

**Supplemental Figure 2:**
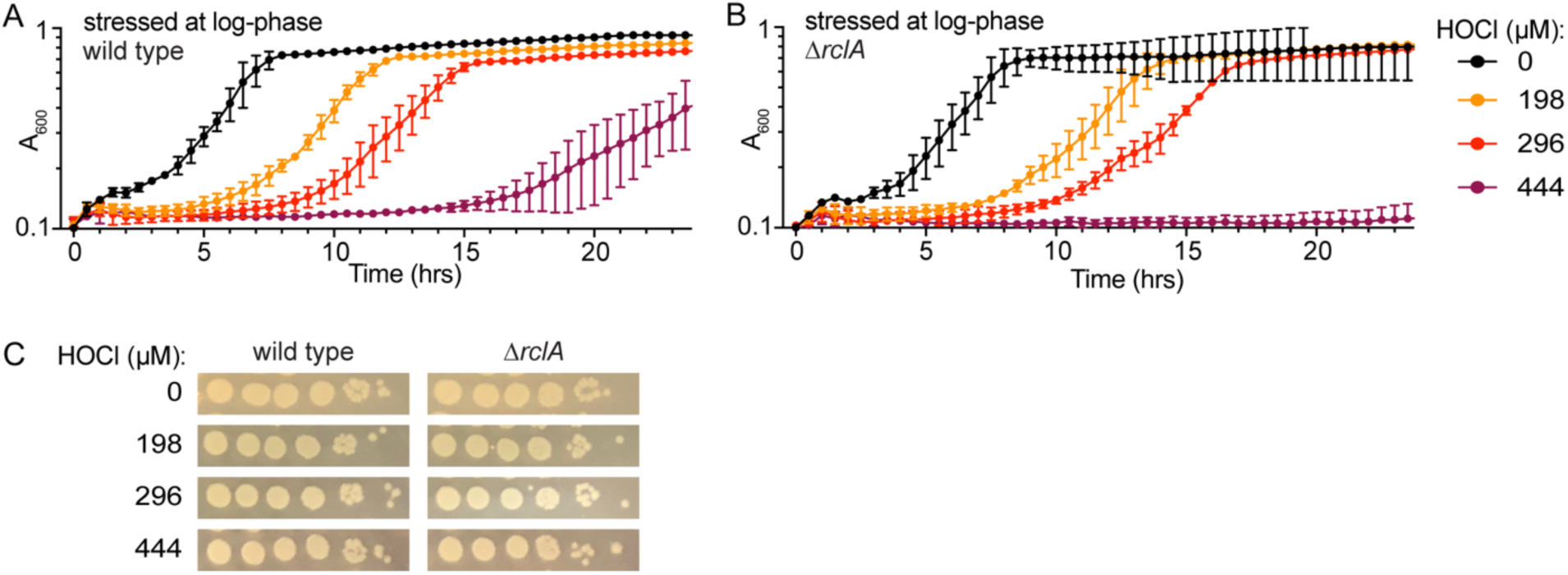
Lag phase extension of HOCl-treated cells in logarithmic growth phase. HOCl growth curves were performed as in Supplemental Figure 1 but on log-phase cultures of each strain. To prepare the log-phase cultures, saturated overnight cultures were subcultured in MOPS glucose and grown up at 37 °C with shaking at 200 rpm until A_600_ = 0.5. *E. coli* lacking *rclA* (B) had increased lag-phase extensions (LPE) relative to wild type (A) at all the tested HOCl concentrations. The observed growth delays of the strains in the presence of HOCl did not correspond with killing, as determined by viability spot plates (C, pictures representative of 3 biological replicates). Viability spot plates were prepared by collecting 5 µL aliquots from each well used for growth curves 10 minutes after adding MOPS ± HOCl to the OD-normalized cells. Serial dilutions in PBS of the aliquots were performed and 5 µL of the dilutions were spotted on LB plates and grown overnight at 37 °C.

**Supplemental Figure 3:**
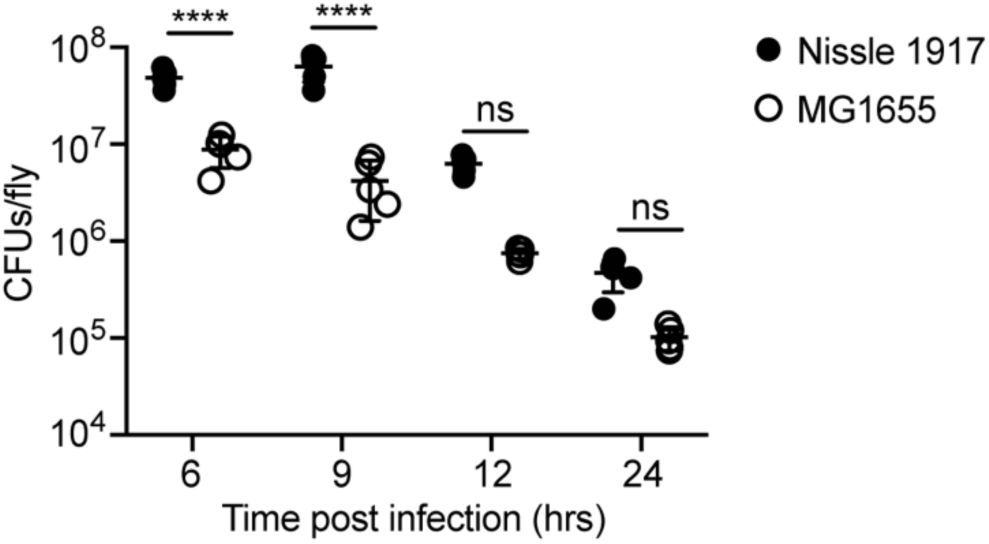
EcN colonizes Canton^S^ *D. melanogaster* to a greater extent than MG1655. Flies were fed either *E. coli* MG1655 or EcN and bacterial loads were measured (n = 5, ± SD). Statistical analysis was performed using a two-way ANOVA with Tukey’s multiple comparisons test (**** = P < 0.0001, ns = not significant).

**Supplemental Figure 4:**
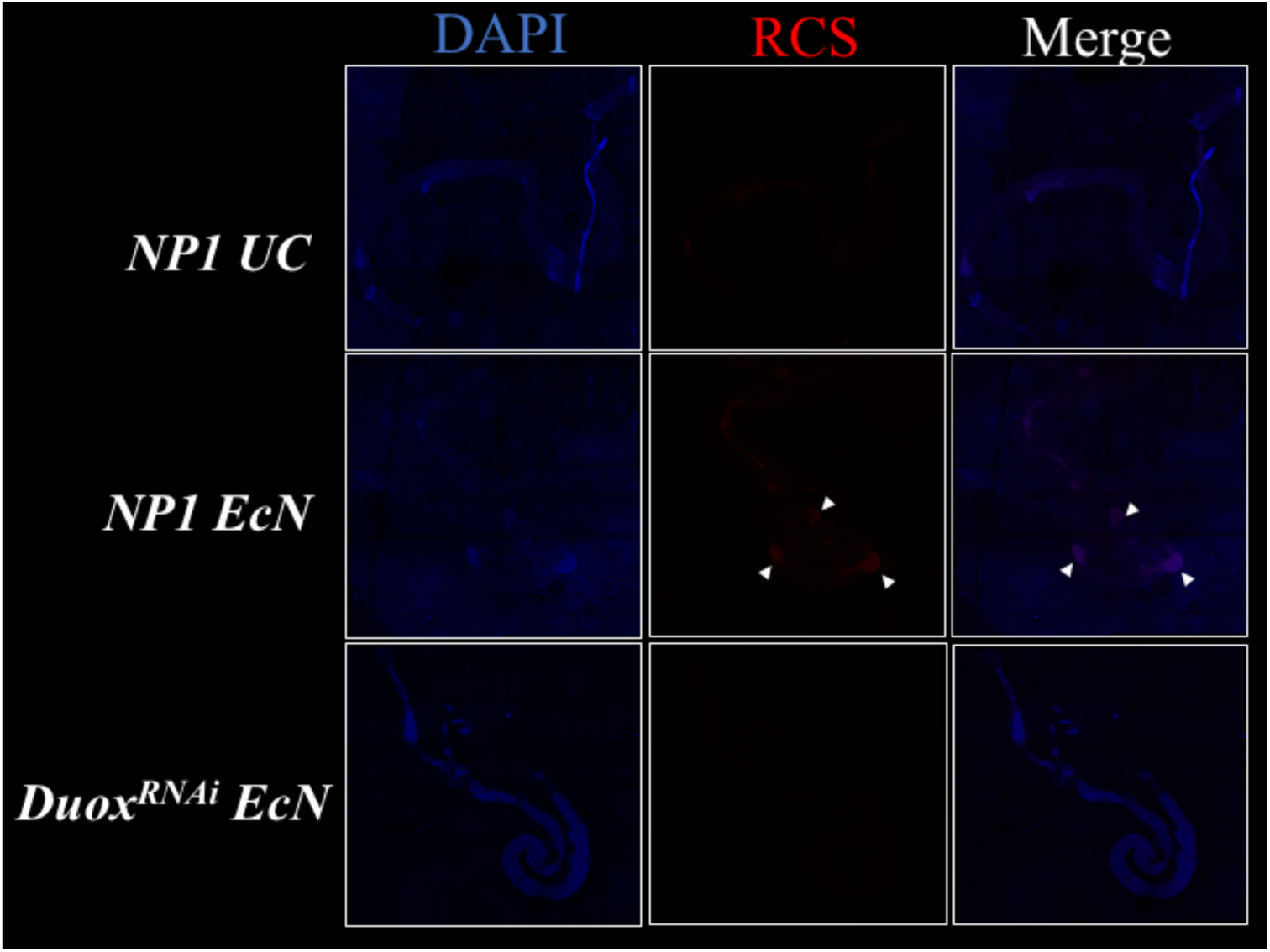
Host-produced HOCl production is silenced by Duox RNAi. *Canton-S* female flies were starved for 2 hours at 29°C. They were then administered a 1:1 mixture of either *E. coli* Nissle (OD_600_=200) or LB and 100 µM R19-S prepared in 5% sucrose. After 30 minutes of feeding, all treatments were switched to sucrose/R19-S mixture. 90 minutes after the initial infection, guts were dissected in PBS and fixed in 4% formaldehyde for 50 minutes. Guts were mounted in Vectashield/DAPI and visualized on a Leica SP8 confocal microscope with a red emission range of 530-603.

**Supplemental Figure 5:**
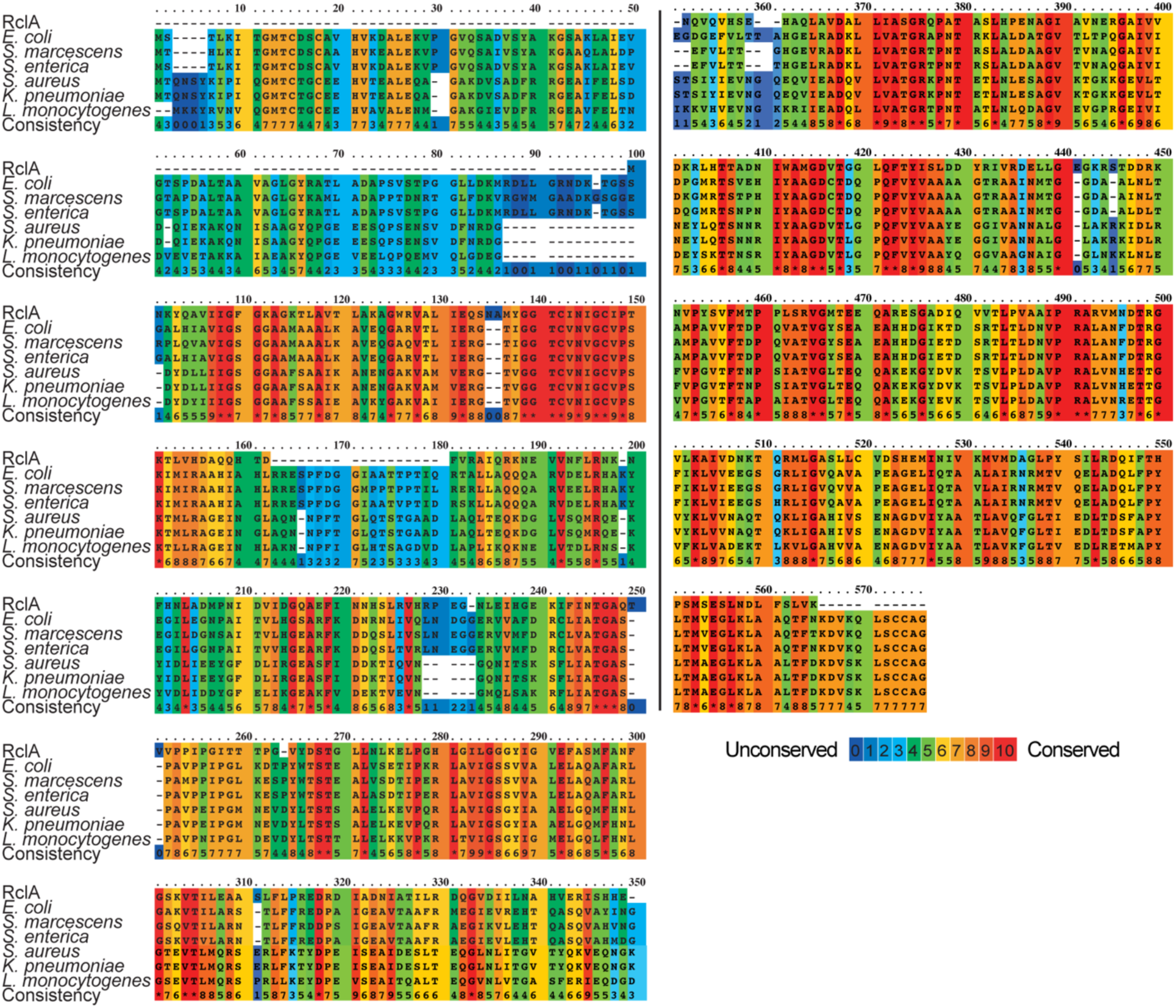
Alignment of full length RclA with MerA sequences from representative bacterial species. Alignment of *E. coli* RclA (ADC80840.1) and MerA amino acid sequences from seven species (*Escherichia coli*, ADC80840.1; *Staphylococcus aureus*, AKA87329.1; *Salmonella enterica*, ABQ57371.1; *Listeria monocytogenes*, PDA94520.1; *Klebsiella pneumoniae*, ABY75610.1; *Serratia marcescens*, ADM52740.1). Alignment, conservation scoring, and graphic were made using PRALINE.

**Supplemental Figure 6:**
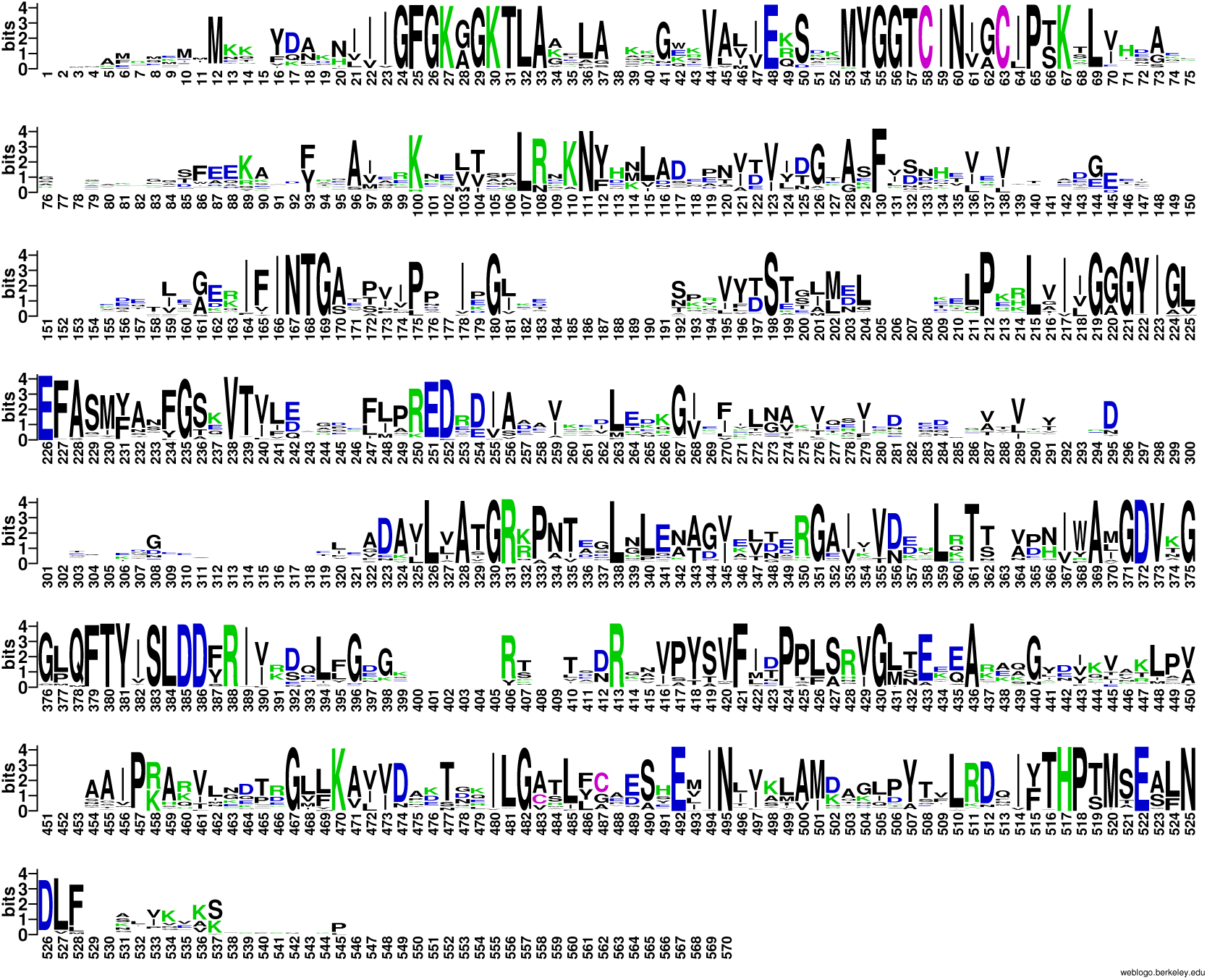
Amino acid conservation in RclA sequences from 284 species. Alignment of RclA sequences made using MUSCLE (BLAST e value < 1×10^-90^ in 284 species) and graphic made using Weblogo (weblogo.berkeley.edu/logo.cgi). Sequences are from the species represented in Figure 1 and listed in Supplemental Data Table 1.

**Supplemental Figure 7:**
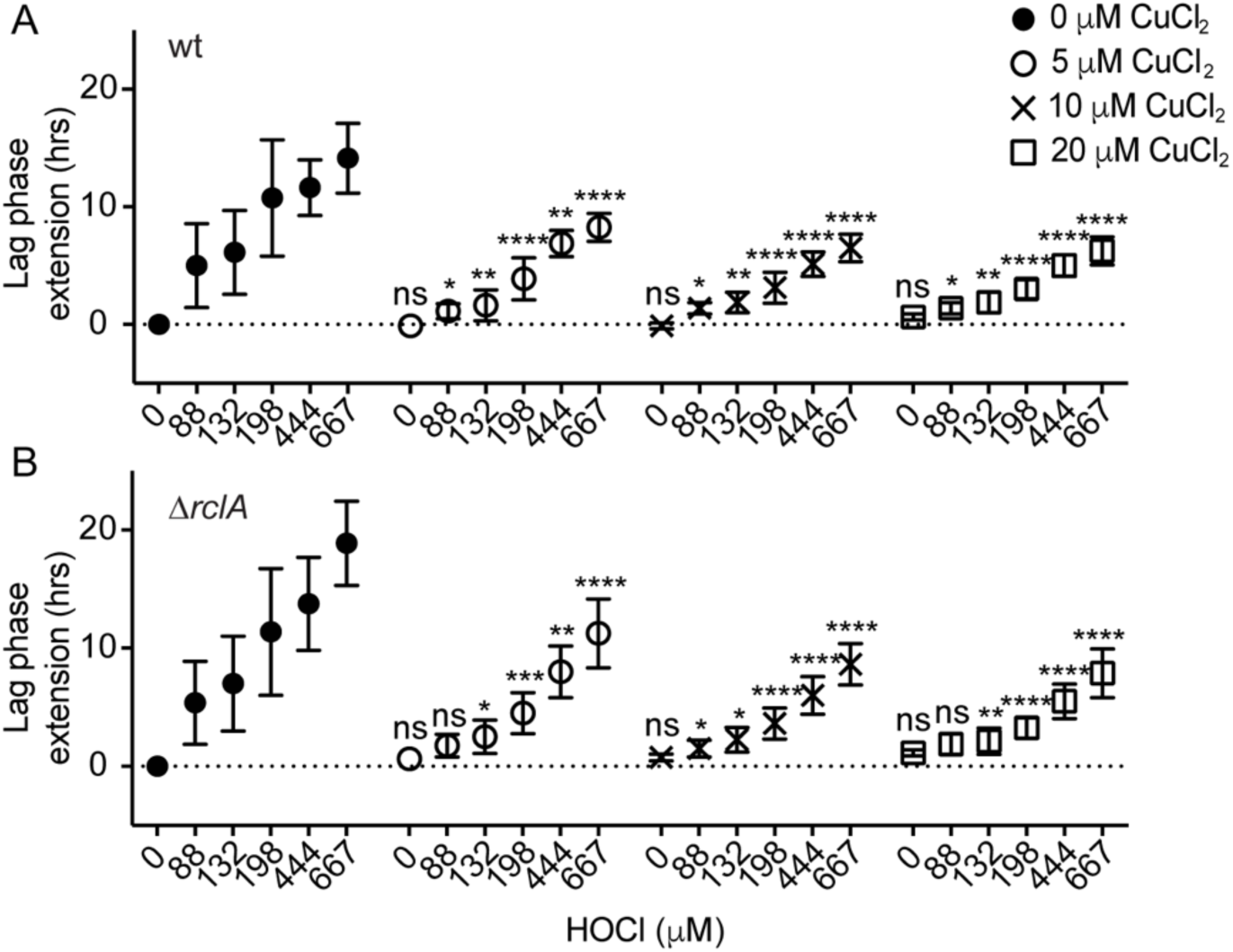
Extracellular CuCl_2_ reduces LPE for both wild-type and Δ*rclA E. coli* grown in the presence of HOCl. LPE determination for wild-type (A) and Δ*rclA* (B) strains grown in the presence of the indicated HOCl and extracellular CuCl_2_ conditions. Growth curves were performed as described in Supplemental Figure 1 but with supplementing the MOPS glucose with the indicated CuCl_2_ concentrations. HOCl and CuCl_2_ were mixed in MOPS minimal media prior to diluting cells A_600_-normalized to 0.08 into each media type. LPEs were calculated by determining the difference in time (hrs) to reach A_600_ > 0.15 for each condition relative to no stress controls. Sensitivity was assessed within strains by comparing LPEs for each HOCl/CuCl_2_ concentration (n = 4, ± SD) at each HOCl concentration (two-way ANOVA for each strain with Dunnett’s multiple comparison using the no CuCl_2_ control for each HOCl concentration, respectively; **** = P < 0.0001, *** = P < 0.001, ** = P < 0.01, * = P < 0.05, ns = not significant).

**Supplemental Figure 8:**
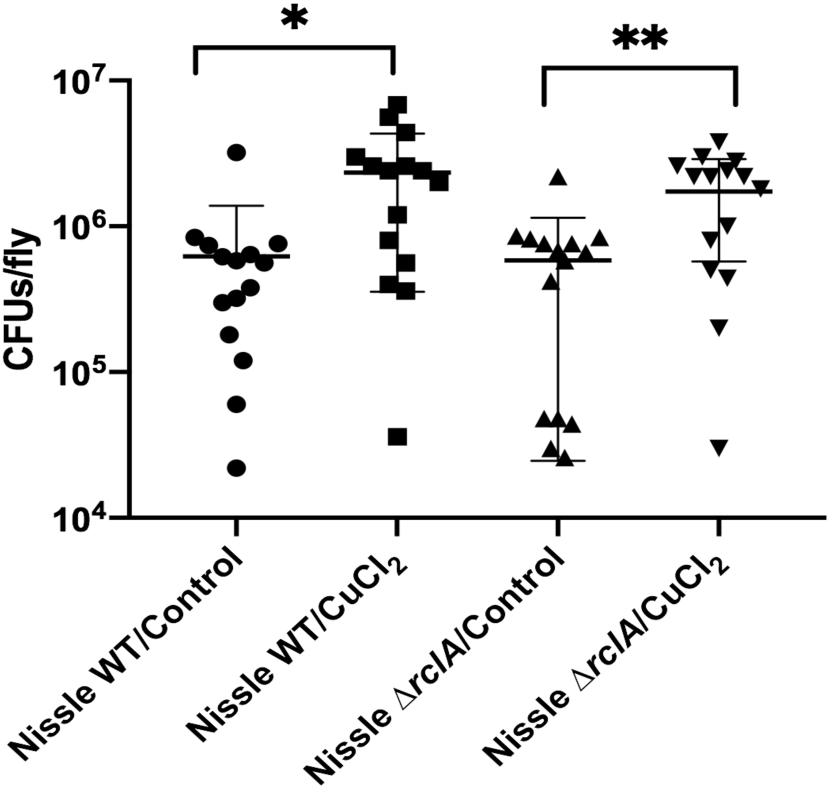
EcN colonization is improved when flies are fed copper. *D. melanogaster* Canton^S^ females fed for 24 hours on a mixture containing sucrose and 1mM CuCl_2_ (or sterile water), which was applied to fly food on a paper disk. Flies were then transferred to new food tubes containing sucrose and either *E. coli* Nissle wild-type (WT) or *E. coli* Nissle Δ*rclA*. Individuals were collected, surface sterilized, homogenized, and plated on LB agar at various time points to determine bacterial load (CFUs/fly). Statistical analysis was performed with GraphPad Prism using a multiple t test with Sidak-Bonferroni correction for multiple comparisons. (N=3, n=60).

**Supplemental Figure 9:**
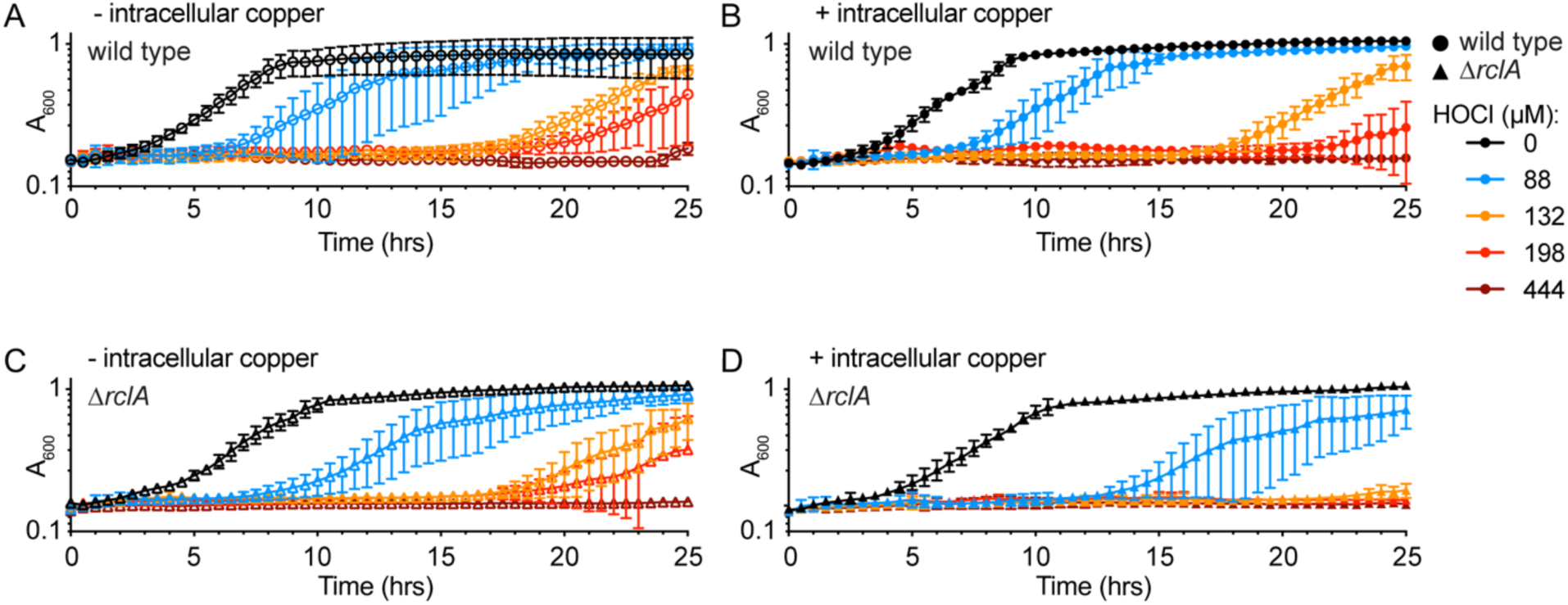
Absence of intracellular copper reduces HOCl sensitivity of *E. coli* lacking *rclA*. Wild type (A and B) and Δ*rclA* (C and D) *E. coli* were grown up overnight in MOPS glucose with (B and D) and without copper (A and C). HOCl growth curves were performed as described in Figure 6.

**Supplemental Figure 10:**
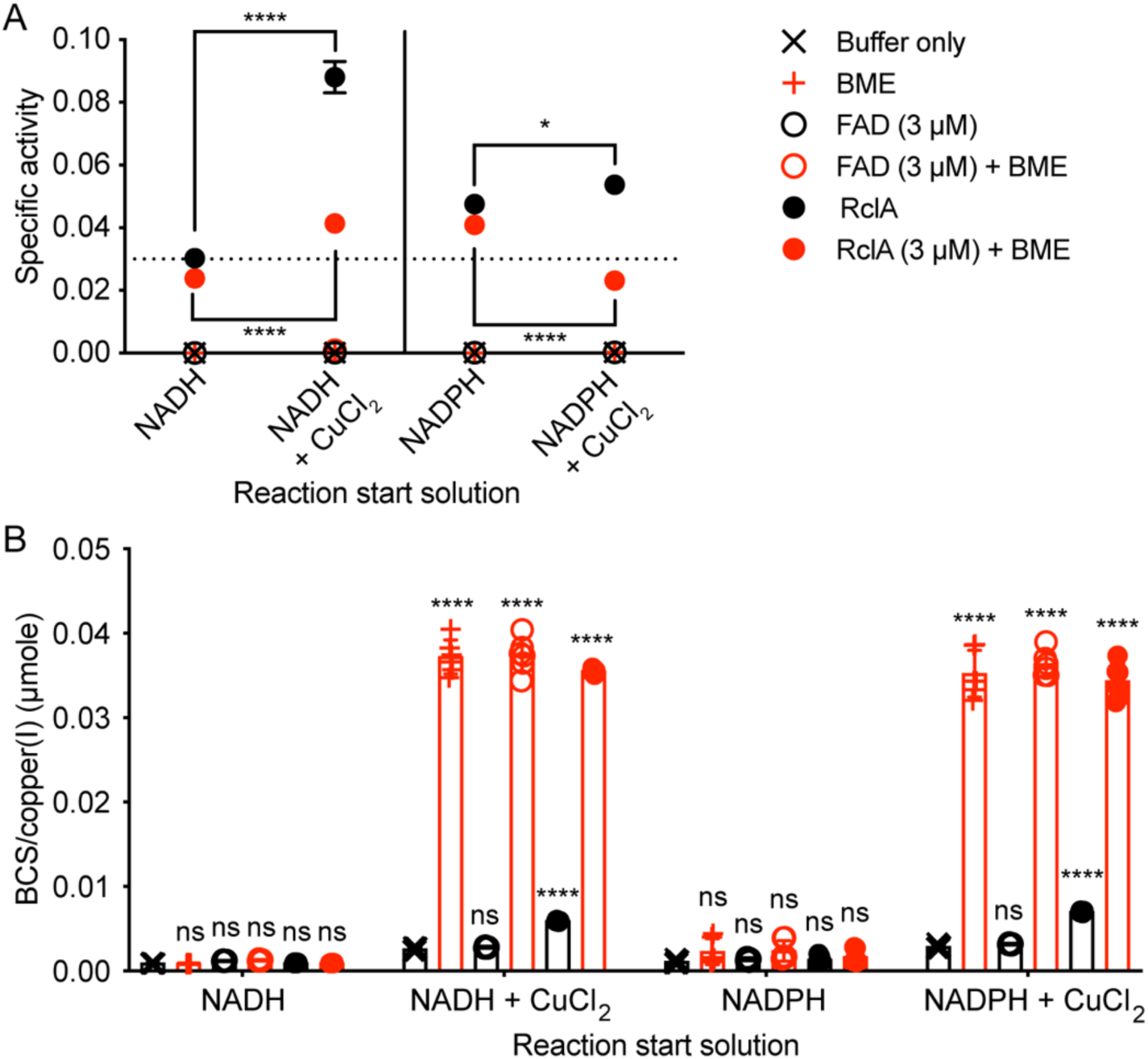
Adding BME to the start solutions of RclA reactions does not increase SA and causes Cu(I) accumulation. RclA reactions performed as in Figure 7 but with the addition of 1 mM BME for indicated samples (n = 6, ± SD). SA (A) and BCS/Cu(I) concentration determined as in Figure 7. Differences in SA in the presence of each metal were analyzed using a two-way ANOVA with Dunnet’s multiple comparison test using the buffer only sample as the control for each start solution (****= P < 0.0001).

**Supplemental Figure 11:**
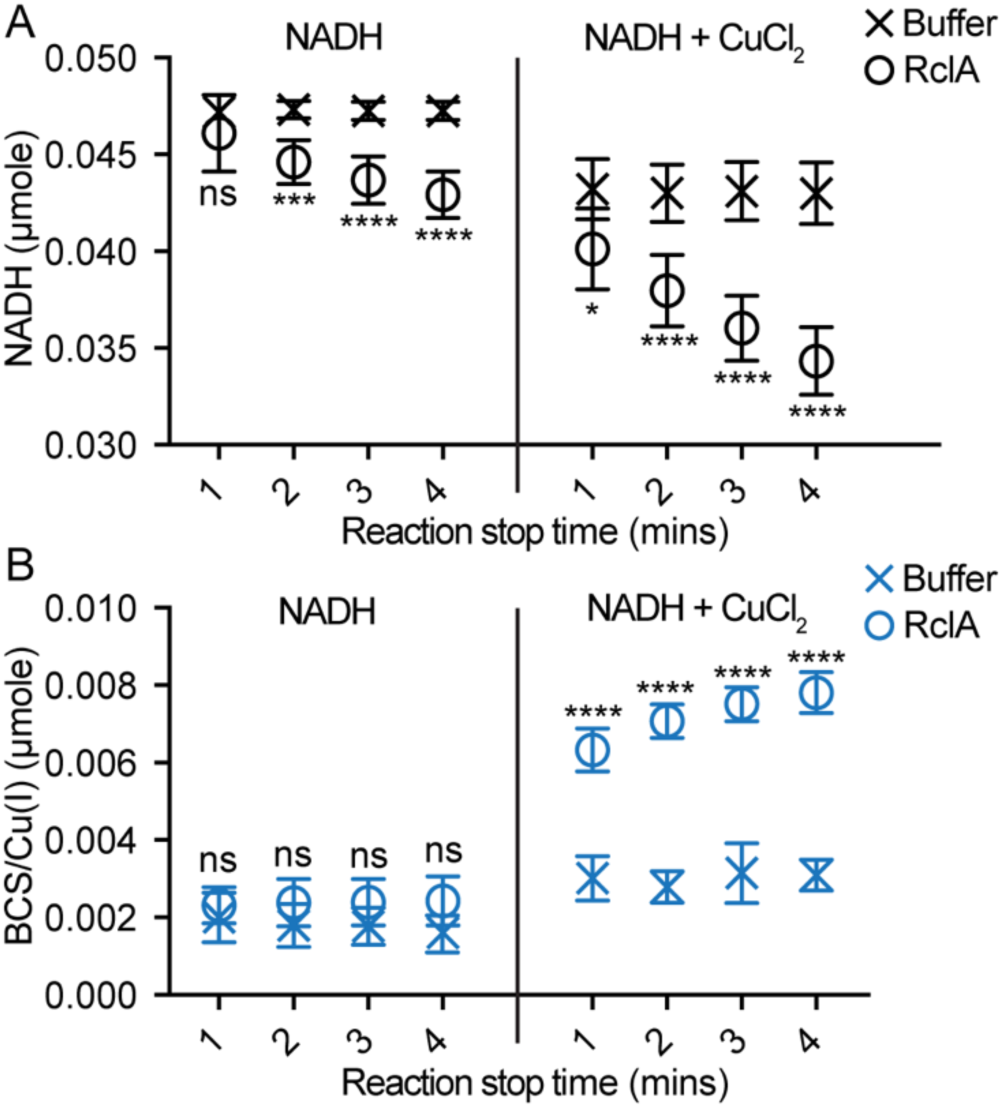
Changes in NADH and copper (I) concentrations over time during the RclA reaction. NADH oxidation (A) and BCS/Cu(I) complex formation (B) were measured over time spectrophotometrically with and without RclA. Reactions were started by adding 100 µL of NADH (200 µM final) or NADH and CuCl_2_ (both 200 µM final) to 100 µL of RclA (3 µM final) at 37 ^°^C using the injector system of a Tecan Infinite M1000 plate reader. All reactions were carried out in 20 mM HEPES, 100 mM NaCl, pH 7. NADH absorbance at 340 nm was measured each minute for the indicated reaction times. Each reaction was then stopped at the indicated times with 10 µL of a BCS (400 µM final) and EDTA (1 mM final) solution using the injector system of the plate reader (n = 6 for each reaction type, ± SD). The stopped reactions were incubated at 37 °C for 5 minutes with the absorbance of BCS/Cu(I) complex being measured at 483 nm each minute to ensure saturation of BCS. Reported BCS/Cu (I) values were calculated using the average of the total absorbance measurements over the five-minute incubation because the values were stable over that time. Differences in the amount of NADH or BCS/Cu(I) complex between the buffer only and RclA reactions were analyzed using a 2-way ANOVA with Sidak’s multiple comparison test (**** = P < 0.0001, *** = P < 0.001, * = P < 0.05, ns = not significant).

**Supplemental Figure 12:**
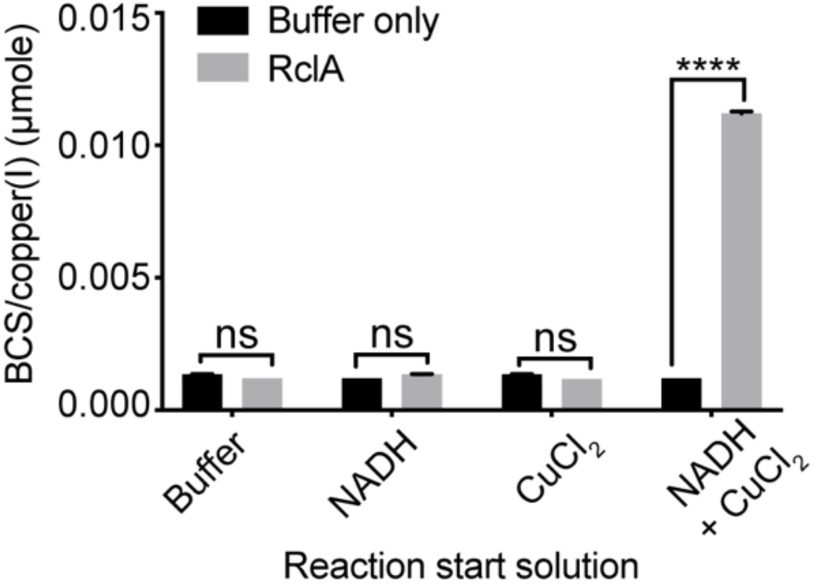
The copper reductase activity of RclA is maintained under anerobic conditions. BCS/Cu(I) complex concentrations were determined as in Figure 7 but all reactions were performed in an anaerobic chamber (n = 6, ± SD). Differences in the amount of BCS/Cu(I) complex between the buffer only and RclA (3 µM) reactions were analyzed using a two-way ANOVA with Tukey’s multiple comparison test (****= P < 0.0001, ns = not significant).

**Supplemental Figure 13:**
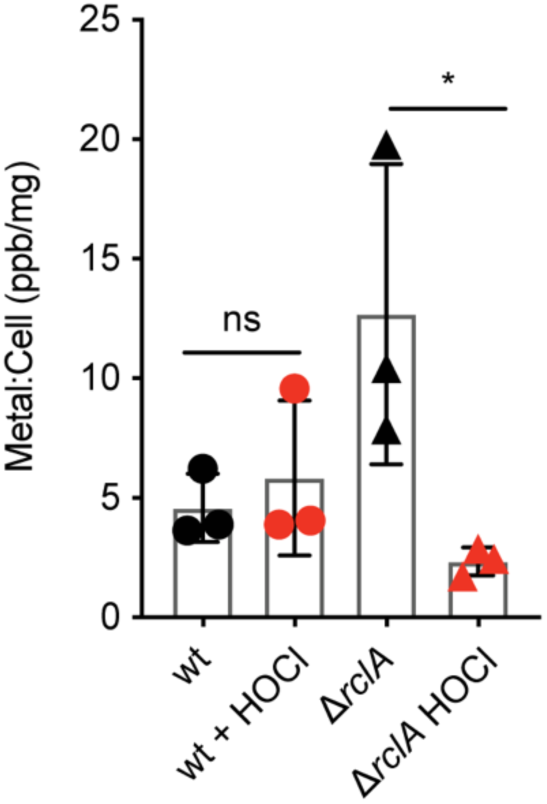
Intracellular copper concentrations do not change after HOCl stress in wildtype *E. coli*. Intracellular copper before and after HOCl stress (30 min at 400 µM) of wild-type (wt) and Δ*rclA* mutant strains was determined using ICP-MS. Differences between intracellular copper analyzed using a two-way ANOVA with Tukey’s multiple comparison test (* = P < 0.05, ns = not significant).

**Supplemental Figure 14:**
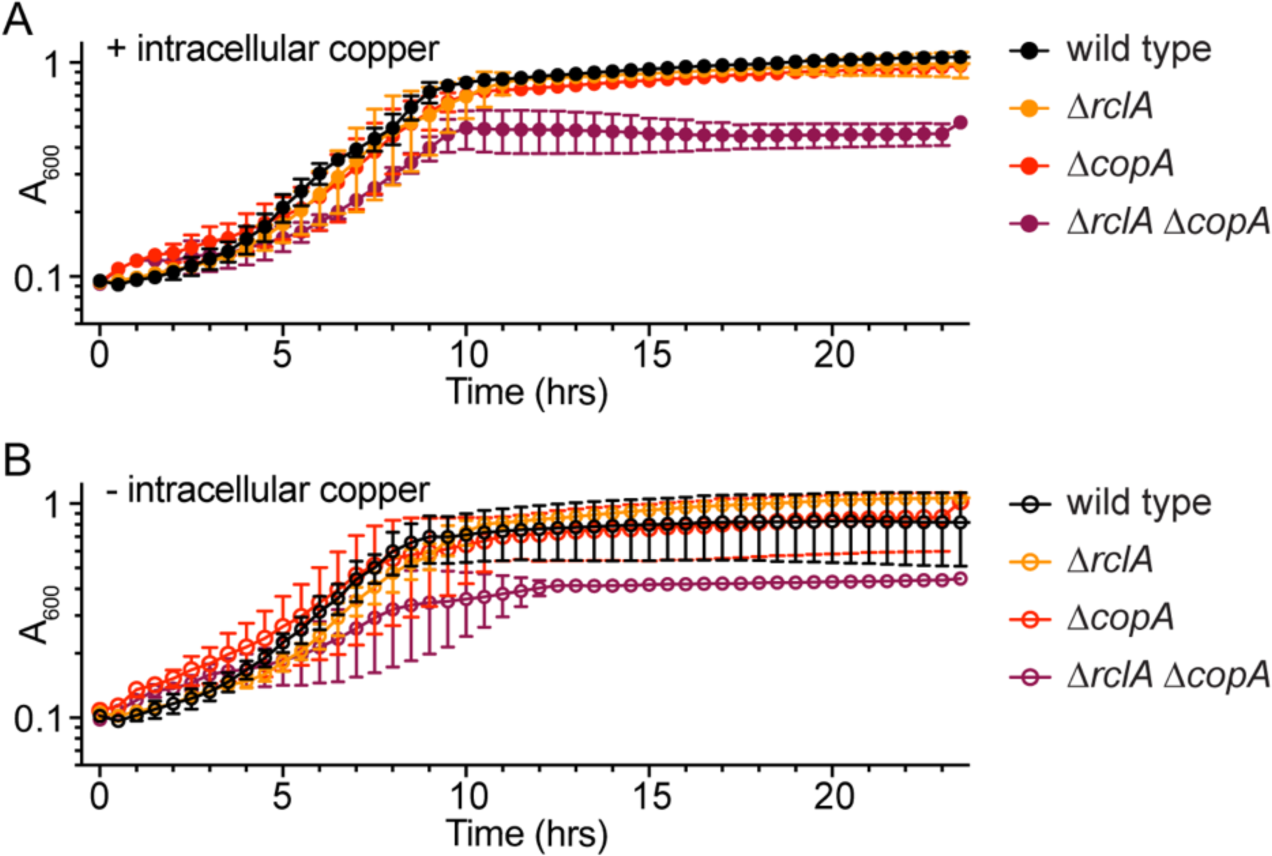
*rclA copA* mutants have a growth defect in copper-free media. A double mutant of *rclA* and *copA* has a growth defect when grown in copper-free MOPS when subcultured from overnight cultures grown either with (A) or without copper (B). HOCl growth curves on cells with and without intracellular copper were performed as described in Figure 6 with copper-free MOPS glucose (n = 3).

**Supplemental Figure 15:**
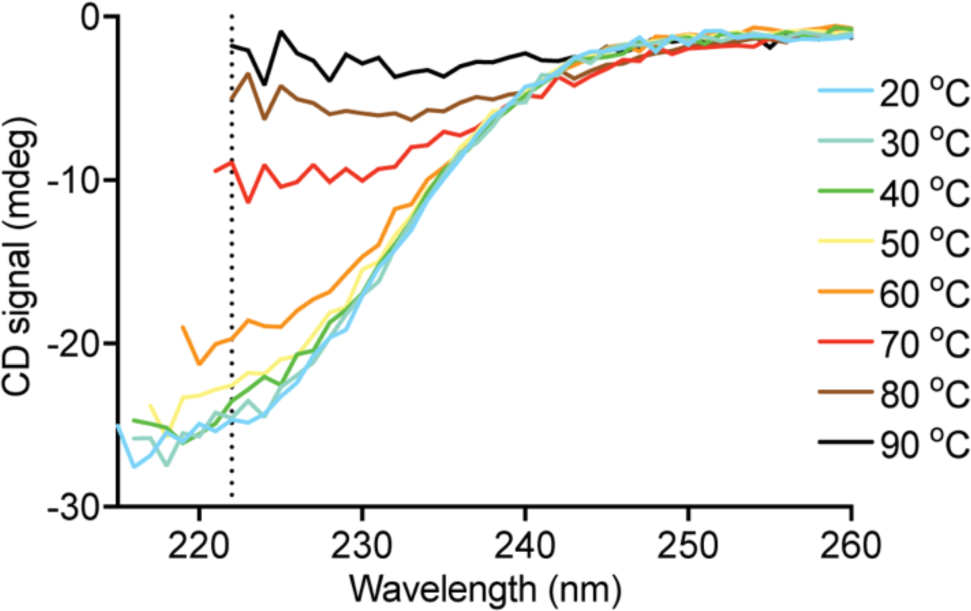
The melting temperature (T_m_) of RclA is approximately 10 °C higher than the average T_m_ of the *E. coli* proteome. (A) CD spectra of RclA measured at indicated temperatures; 222 nm is indicated with a vertical dashed line.

**Supplemental Figure 16:**
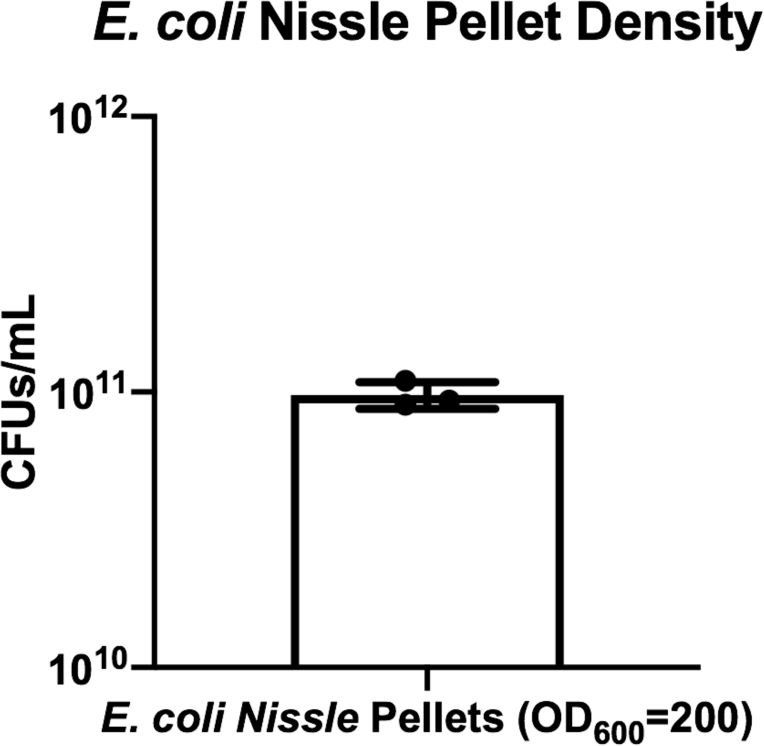
*E. coli Nissle* pellets contain consistent bacterial density. Three independent *E. coli Nissle* cultures were grown in 30 ml of LB at 37°C with shaking for 20 hours. Cultures were pelleted by centrifugation at 3,428 x g. Resulting pellets were adjusted to an OD_600_ of 200. Dilutions were plated to determine CFUs/ml.

